# Natural heteroclitic-like peptides are generated by SARS-CoV-2 mutations

**DOI:** 10.1101/2022.10.28.513849

**Authors:** Camilla Tiezzi, Andrea Vecchi, Marzia Rossi, Davide Cavazzini, Angelo Bolchi, Diletta Laccabue, Luca Sacchelli, Federica Brillo, Tiziana Meschi, Andrea Ticinesi, Antonio Nouvenne, Gaetano Donofrio, Paola Zanelli, Magda Benecchi, Silvia Giuliodori, Paola Fisicaro, Ilaria Montali, Simona Urbani, Giuseppe Pedrazzi, Gabriele Missale, Amalio Telenti, Davide Corti, Simone Ottonello, Carlo Ferrari, Carolina Boni

**Author notes:** These authors contributed equally to this work (Camilla Tiezzi, Andrea Vecchi, Marzia Rossi). These authors jointly supervised this work (Carlo Ferrari, Carolina Boni). **Corresponding authors**: Prof. Carlo Ferrari, Department of Medicine and Surgery, University of Parma, Laboratory of Viral Immunopathology, Unit of Infectious Diseases and Hepatology, Azienda Ospedaliero-Universitaria di Parma, Via Gramsci 14, 43126 Parma, Italy., Dr. Carolina Boni, Laboratory of Viral Immunopathology, Unit of Infectious Diseases and Hepatology, Azienda Ospedaliero-Universitaria di Parma, Via Gramsci 14, 43126 Parma, Italy.

## Abstract

Mutations carried by SARS-CoV-2 spike protein variants may promote viral escape from immune protection. Humoral immunity is sensitive to evasion by SARS-CoV-2 mutants, but the impact of viral evolution on the interplay between virus and host CD8 T cell reactivity remains uncertain. By a systematic functional analysis of 30 spike variant mutations, we show that in vaccinated as well as convalescent subjects, mutated epitopes can have not only a neutral or abrogating effect on the recognition by CD8 T cells but can also enhance or even generate de novo CD8 T cell responses. Large pools of peptides spanning the entire spike sequence and comprising previously identified CD8 T cell epitopes were then used in parallel with variant peptides to define strength and multispecificity of total anti-spike CD8 responses. In some individuals, CD8 cells were narrowly focused on a few epitopes indicating that in this context of weak and oligospecific responses the overall antiviral protection can likely benefit of the function enhancing effect of heteroclitic-like mutations. In conclusion, appearance of mutated stimulatory epitopes likely reflects an epiphenomenon of SARS-CoV-2 evolution driven by antibody evasion and increased transmissibility, that might bear clinical relevance in a subset of individuals with weak and oligospecific CD8 T cell responses.

## INTRODUCTION

RNA viruses tend to accumulate mutations more rapidly and extensively than DNA-based pathogens^1,2^. The main cause of this enhanced mutability is low-fidelity replication, which leads to genetically diverse but related virus variants populations. Such variants may progressively acquire better tropism as well as higher infectivity and pathogenicity, with possible acquisition of resistance to vaccines and antiviral compounds. As an RNA virus, evolution of the SARS-CoV-2 genome resulted in multiple variants (so-called ‘Variants of Concern’, VoC and ‘Variants of Interest’ VoI) that have progressively acquired an increased transmission capacity^3–12^. Understanding the effect of mutations on protective immune responses is key to the development of new generation vaccines and antivirals.

As a general rule, new mutations tend to be fixed either because of the acquisition of an enhanced transmissibility or because of escape from immunological pressure. Both mechanisms are thought to be causally involved in the selection of SARS-CoV-2 variants. In the absence of pre-existing immunity, as in the case of the early SARS-CoV-2 outbreak, escape from immune responses is expected to contribute very little to the rate of virus spread^13^, which should be primarily determined by a direct effect of mutations on virus tropism, efficiency of replication and infection. However, as protective immunity becomes more widespread, escape from neutralizing antibodies and cytotoxic CD8 T cell responses can provide a more relevant contribution to virus transmissibility. Indeed, the establishment of new variants, such as beta, gamma, delta and omicron, has been associated with a progressively growing impact of virus mutations on neutralizing antibody responses^14–19^. Remarkably, the impact of SARS-CoV-2 variability seems to be instead much lower on CD8 T-cell-based immunity^14,15,20–25^. This differential effect of mutations is in keeping with the significant rate of breakthrough infections by the delta and omicron variants in Wuhan type-based spike vaccinated individuals, in which, however, severe symptomatic disease is dampened possibly through the contribution of CD8 T cell-mediated immunity ^26,27^.

Considering the important role of CD8 T cell responses in antiviral protection^28–31^, we performed a comprehensive investigation of the interplay between SARS-CoV-2 variation and spike vaccine-induced CD8 T cells reactivity. Strikingly, we discovered an unexpectedly enhanced immune responsiveness to a specific subset of mutated spike CD8 T cell epitopes. The possibility of enhancing the immune stimulatory activity of previously identified CD8 T cell epitopes by single amino acid changes, with generation of variant peptides named heteroclitic, has widely been explored in vitro, as a possible strategy to improve the antiviral protection of T cell-based vaccines and therapies. Instead, here we report the natural generation of variant peptides able to ameliorate responsiveness to the corresponding prototype sequences contained in the Wuhan-based vaccine. These findings offer novel perspectives on the breadth of protection that may be conferred by CD8 T cells and on their role in the evolutionary biology of SARS CoV-2.

## RESULTS

### Impact of variant mutations on SARS-CoV-2-specific T cell responses

To examine the effect of SARS-CoV-2 variant mutations on CD8 T cell reactivity, we studied 36 vaccinated donors, naïve to infection, 1-2 months after the second dose of the Pfizer/BioNTech vaccine. All subjects showed high levels of serum anti-spike neutralizing antibodies detected by a SARS-CoV-2 Wuhan-Hu-1 pseudovirus neutralization assay (Extended Data Table 1). To study the CD8 T cell response to SARS-CoV-2 spike (S), we used an ex-vivo FluoroSPOT assay based on PBMC stimulation with 10 mer peptides (overlapping by 9 residues) spanning VOC- and VOI-containing S regions harboring all the main mutations that emerged since the initial appearance of SARS-CoV-2 in Wuhan until June 2021. For each S mutation, two peptide pools were synthesized in order to cover both the Wuhan prototype and the subsequently mutated SARS-CoV-2 variants and to accommodate each individual mutation at each position of the overlapping peptides (Extended Data Tables 2 and 3). The ability of each peptide pool to simultaneously elicit IFN-γ, TNF-α and IL-2 production by CD8 T cells was then measured and stringent criteria were applied in order to identify statistically significant SARS-CoV-2 spike-specific T cell responses (see ‘Methods’ for details).

SARS-CoV-2 spike-specific CD8 T cell responses were detected against at least 1 peptide pool in 17 out of the 36 tested vaccinees (Fig. 1). All peptide pools were able to stimulate significant T cell responses in at least one responder individual, showing that each tested mutation was contained within CD8 T cell epitopes. As expected, some mutations had a neutral effect, meaning that prototype (*grey bars*) and mutated (*black bars*) peptides were similarly capable of stimulating CD8 T cell responses. Other mutations negatively affected T cell responses because the variant peptide exhibited a reduced stimulatory activity compared to the corresponding prototype sequence. Surprisingly, however, the opposite effect was observed with a subset of mutations, which conferred to the corresponding variant peptides a significantly enhanced stimulatory potency compared to the original prototype peptide.

**Fig. 1.**
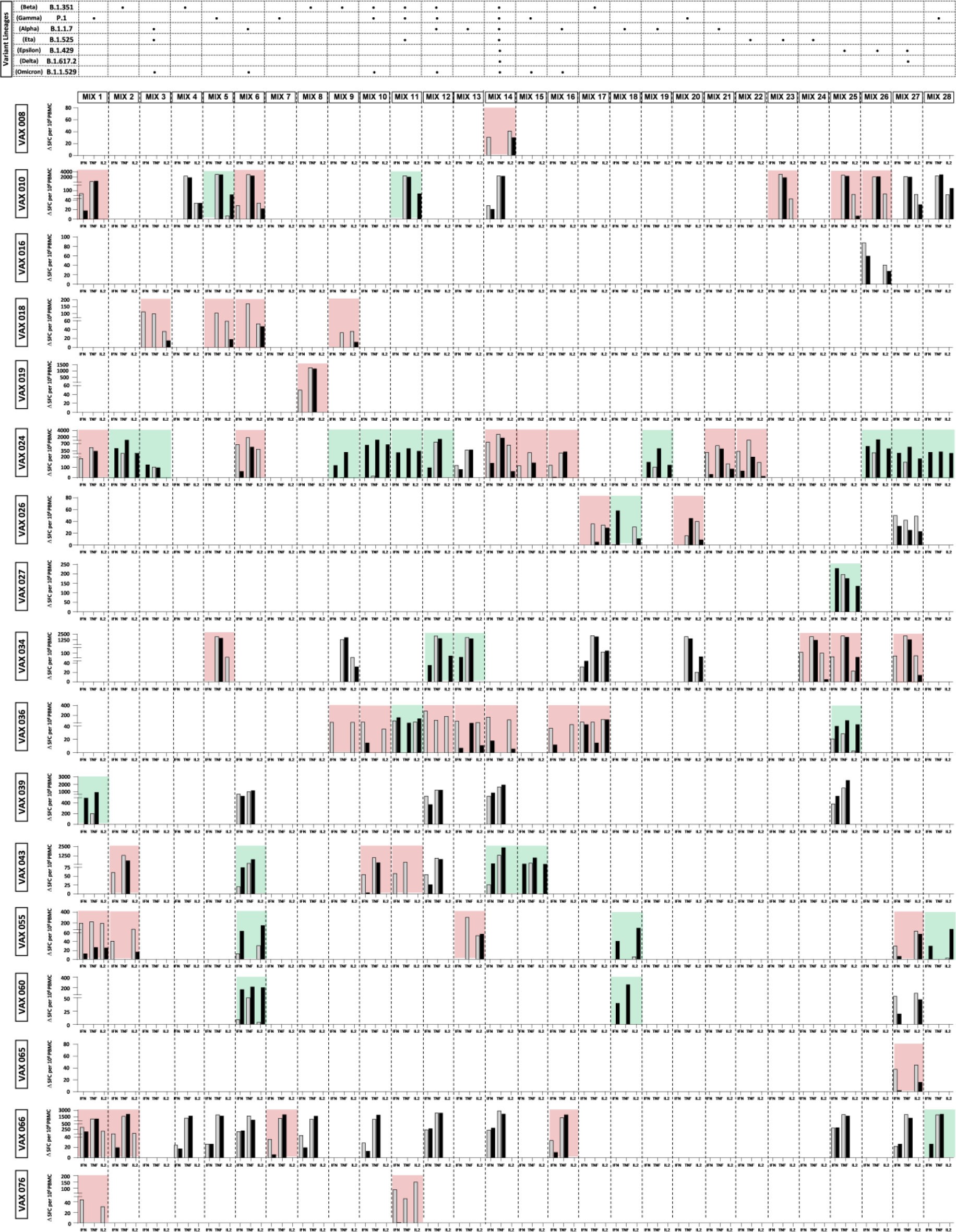
Impact of VOC and VOI mutations on spike-specific T cell responses measured in BNT162b2 vaccinated donors. PBMCs from vaccinated individuals (n=36) were stimulated for 18 hours with overlapping peptides covering the spike sequences containing VOC and VOI mutations as well as the corresponding sequences of the prototype Wuhan SARS-CoV-2 strain. IFN-γ, TNF-α and IL-2 secreting T cells were measured by FluoroSPOT assay (spot-forming cells/SFCs per 10^6^ PBMCs). Grey and black bars represent the frequency of IFN-γ-, TNF-α- or IL-2-SFCs reactive to individual spike peptide pools spanning either the wild type or the VOC and VOI mutated strains, respectively. Red- and green-shadowed areas indicate significantly diminished or increased T cell responses induced by mutated compared to wild type peptide epitopes, respectively; unshadowed (*white background*) areas represent neutral responses. The location of VOC and VOI mutations within each peptide pool is indicated at the top of the figure. Spike sequence positions of each peptide pool are reported in Extended Data Table 2. FluoroSPOT results were considered positive if the number of spots in the stimulated wells was at least three standard deviations above background and the difference between the number of spots in the stimulated and unstimulated wells was above 10. Only vaccinated subjects that positively responded to at least two out of the three tested antiviral cytokines (17 out of 36 individuals) are shown. (B.1.351, Beta; P.1, Gamma; B.1.1.7, Alpha; B.1525, Eta; B.1.429, Epsilon; B.1.617.2, Delta; B.1.1.529, Omicron).

To unambiguously classify enhancing and inhibitory effects of individual mutations, we calculated the fold-change for each T cell response stimulated by a mutant peptide pool relative to the corresponding unmutated pool (mutated/wild-type) and then applied an *effect size* filter to quantitatively evaluate the relationship between variables (see ‘Methods’ for details). Keeping the *large effect* threshold as a discriminant, only T cell responses below or above the *large effect size* values were considered as significantly decreased or increased (< − 0.48 or > + 0.48, respectively; see Extended Data Fig. 1A). In addition, to provide an even more stringent definition of the modulatory effects caused by individual mutations, only responses featuring a concordant modulation of at least two of the three tested cytokines were considered as reliable indicators of a positive or a negative effect.

In some cases, negative or positive effects were clearly appreciable even without statistical analysis, because the mutated peptides either induced a response not observed with the prototype sequence, or failed to induce any detectable response as observed with the corresponding prototype peptide (in these cases only a *grey* or a *black* bar is displayed in Fig. 1). Overall, a statistically supported neutral effect was detected in 25% of the responses (Fig. 1 and Extended Data Fig. 1). A markedly decreased or no response to mutated spike variant peptide pools was observed and statistically validated in 45 cases (46% of the total responses), distributed among 13 of the 17 responder vaccinated individuals (76%) (Fig. 1 and Extended Data Fig. 1A). In contrast, and quite surprisingly, the opposite phenomenon, i.e., a significantly enhanced response induced by a subset of mutated spike peptides, was observed in 11 of the 17 vaccinated responders (64%), accounting for 29% of all significant responses induced by peptide pool stimulation (Fig. 1 and Extended Data Fig. 1A). As further shown in Extended Data Fig. 1B (*top panel*), enhanced responses to variant peptides were particularly evident for IFN-γ and IL-2 production.

Moreover, as revealed by simultaneous detection of IFN-γ, TNF-α and IL-2 at a single T cell level, enhanced cytokine production was sustained not only by an increased frequency of the overall population of cells producing individual cytokines upon stimulation (sum of single, double and triple positive cells) but also by polyfunctional T cells capable of producing IFN-γ, TNF-α and IL-2 simultaneously (Extended Data Fig. 2).

The enhancing or inhibitory effects associated with individual mutations detected in the initial screening (Fig. 1) were confirmed by dose titration experiments using different concentrations of wild-type and variant peptide pools (Fig. 2A and Extended Data Fig. 3A). For example, a clear dose-dependent response was obtained with wild-type MIX 1 tested on cells from VAX 055, whereas a barely detectable response was induced by the corresponding mutated sequences. In contrast, mutated MIX 6 induced much stronger responses in VAX 055 than the corresponding wild-type pool. Of note, the varying steepness of the dose-response curves obtained with different vaccinees is likely explained by quantitative differences in peptide affinity recognition by T cells from different vaccinated subjects (Fig. 2A). Specifically, while for certain pools, peptide concentrations as low as 0.1-0.5 μM were sufficient to induce maximal cytokine production (e.g., VAX 055-MIX 1), concentrations of at least 1 μM were required for other peptides (e.g., VAX 036-MIX 12; VAX 076-MIX 11; VAX 055-MIX 6; VAX 036-MIX 11; VAX 060-MIX 6).

**Fig. 2.**
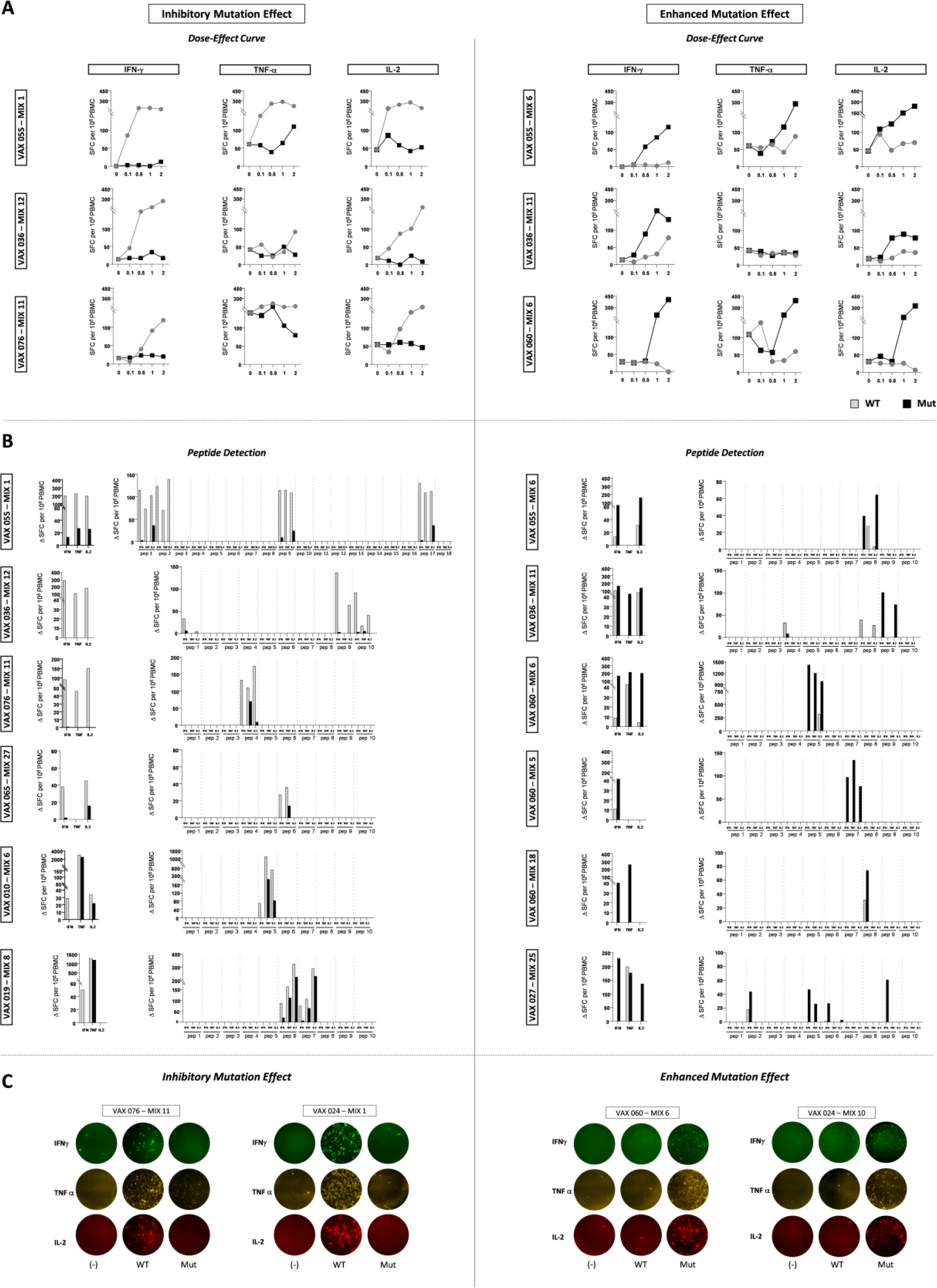
Dose-dependent response to wild-type and variant peptides and identification of individual T cell epitopes. **A)** Dose response curves to the different peptide pools determined by ex-vivo FluoroSPOT using the indicated increasing concentrations of wild-type and variant peptide pools. Grey and black lines and symbols represent T cell responses induced by wild-type (WT) and mutated peptide pools, respectively. **B)** Responses to most of the pools were dissected by using individual component peptides. Individual epitopes were identified by ex-vivo FluoroSPOT assays performed on PBMCs from vaccinated subjects (n=8). Grey and black bars represent the responses induced by wild-type and mutated peptides, respectively. **C)** Representative FluoroSPOT assay results with peptides containing inhibitory or enhancing mutations.

### Epitope identification and characterization of the responder T cell subset

To dissect the responses elicited by the different peptide pools and map the specific amino acid sequences responsible for T cell response induction, individual peptides from pools inducing significant T cell responses were tested for their PBMC stimulation capacity by ex-vivo FluoroSPOT (Fig. 2B). This analysis allowed to deconvolute the individual epitopes within most of the tested pools (8 out of 9). All individual peptides reproduced the same enhancing or inhibitory effect displayed by the corresponding peptide pool. Representative examples of the responses elicited by individual peptides and by the corresponding peptide pools are shown in Fig. 2C (see Extended Data Fig. 3B for a larger set of FluoroSPOT assay data).

The above results, obtained with different concentrations of overlapping peptide pools and individual peptides of optimal length for CD8 T cell recognition, confirmed that while some naturally acquired SARS-CoV-2 spike mutations have either no effect or negatively influence epitope immunogenicity, a subset of mutated epitopes is endowed with an enhanced T cell activation capacity.

Additional experiments were then performed by ex-vivo intracellular cytokine staining (ICS) to further verify FluoroSPOT assay results and to better define the specific T cell sub-populations targeted by SARS-CoV-2 spike peptides. To this end, PBMCs from FluoroSPOT-positive vaccinated donors were stimulated by overnight incubation with mutated or wild-type peptide pools, followed by flow-cytometric measurement of IFN-γ, TNF-α and IL-2 production as well as CD107a degranulation. As suggested by the length of the peptides utilized for this analysis, antiviral cytokine production and CD107 degranulation were entirely sustained by SARS-CoV-2-specific CD8 T cells (Fig. 3A). As shown in Fig. 3B, FluoroSPOT and ICS results were fully concordant. Parallel measurements of CD107 degranulation also gave largely concordant results with only a few exceptions; namely, VAX 060/MIX 6, VAX 066/MIX 1 and VAX 076/MIX 1 (Fig. 3B) where a discordance between cytokine production and degranulation was observed.

**Fig. 3.**
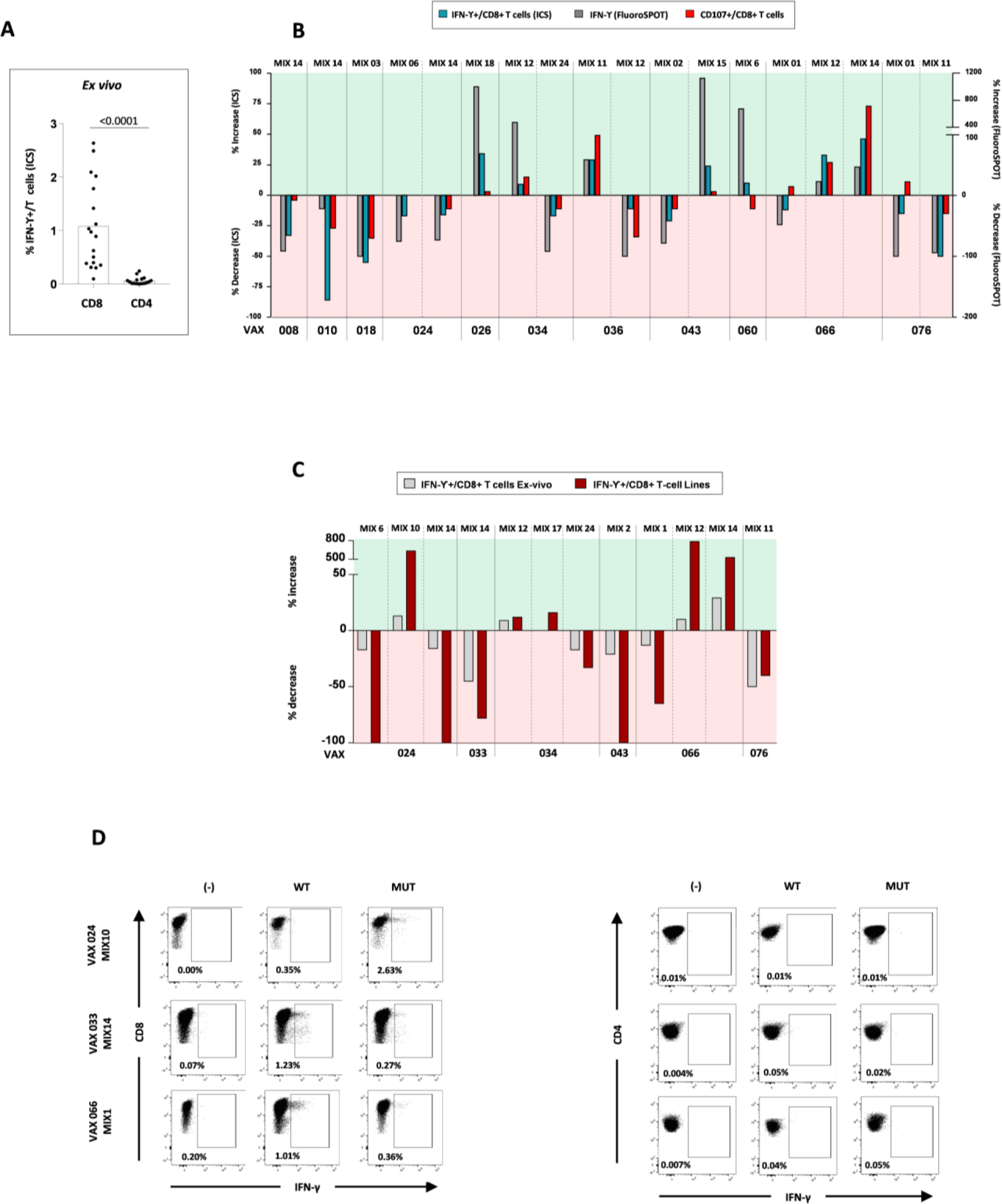
Comparative ex-vivo and in vitro analysis of the variant peptide effect on T cells. **A)** Dots represent the frequency of spike-specific IFN-g-producing CD8 and CD4 T cells reactive to wild-type and variant spike peptide pools determined by flow-cytometry after 18 hours stimulation; background-subtracted data were analyzed statistically by the Mann-Whitney test. **B)** Bars represent decreased or increased (*red-* and *green-*shadowed areas, respectively) responses to variant versus wild-type peptide pools measured by IFN-g production (FluoroSPOT and ICS assays) and CD107 degranulation in 11 vaccinated subjects. **C)** Enhancing and inhibitory effects of mutated versus wild-type peptides measured by ICS on CD8 T cells ex-vivo (*grey bars*) or on in vitro expanded C8 T cell lines (*red bars*); T-cell lines were generated by wild-type and variant peptide pool stimulation for 8 days of PBMCs from six vaccinated donors. **D)**Representative dot-plots derived from cytofluorimetric analysis of in vitro expanded T cells, confirming that the stimulatory and inhibitory effects of variant peptides on IFN-γ production are totally sustained by CD8 T cells.

The enhancing or inhibitory effects detected by ex-vivo FluoroSPOT and ICS assays were reproduced by T cell lines generated upon 8-10 days stimulation of PBMCs with wild-type and mutated peptide pools (Fig. 3C). In keeping with ex-vivo results, responses measured after in vitro T cell expansion were exclusively mediated by CD8 cells, with no detectable contribution by CD4 T cells (Fig. 3D).

### Location of the CD8 response-modulating mutations on the spike protein

To gain insight into the possible structural/functional implications of the modulatory SARS-CoV-2 mutations contained within the identified CD8 epitopes (listed in Fig. 4A), we mapped these epitopes on the three-dimensional structure of the spike protein. As shown in Fig. 4B, nearly all mutated epitopes (20 out of 21) are located within the receptor binding (RBD) and the N-terminal (NTD) domains of the spike S1 subunit, with no apparent distinction between inhibitory and enhancing epitopes. The NTD has been shown to be involved in co-receptor recognition, while the RBD, and particularly the receptor binding motif (RBM), is directly involved in ACE2 receptor engagement. Seven of the 12 inhibitory mutations are located in the NTD, while the remaining five mutations map to the RBM. The latter mutations include the L452R substitution that is shared by various viral lineages (including the currently circulating BA.5 Omicron sublineage) and has been shown to increase viral escape by lowering the affinity of several neutralizing antibodies (nAbs)^32,33^, and the N501Y mutation that strikingly potentiates the RBM-ACE2 interaction^34^. The same number of enhancing mutations is associated to the NTD, but only one is located within the RBM and an additional one (T716I) maps to a β-sheet of the S2 spike subunit that functions as a pivot point during the pre- to post-fusion transition. Three of the seven enhancing NTD-associated mutations, including the Y144 deletion that in different vaccinated subjects was found to elicit both enhancing and inhibitory effects, map to the protruding and highly immunogenic “NTD supersite”, an epitope-rich region targeted by several nAbs^35^. The only RBM-located, enhancing mutation (E484K), which is also shared by different viral lineages, encompasses two epitopes (one inhibitory and the other one enhancing according to our analysis; see Fig. 4B) that are key recognition sites by neutralizing (especially class 2) antibodies^36^.

**Fig. 4.**
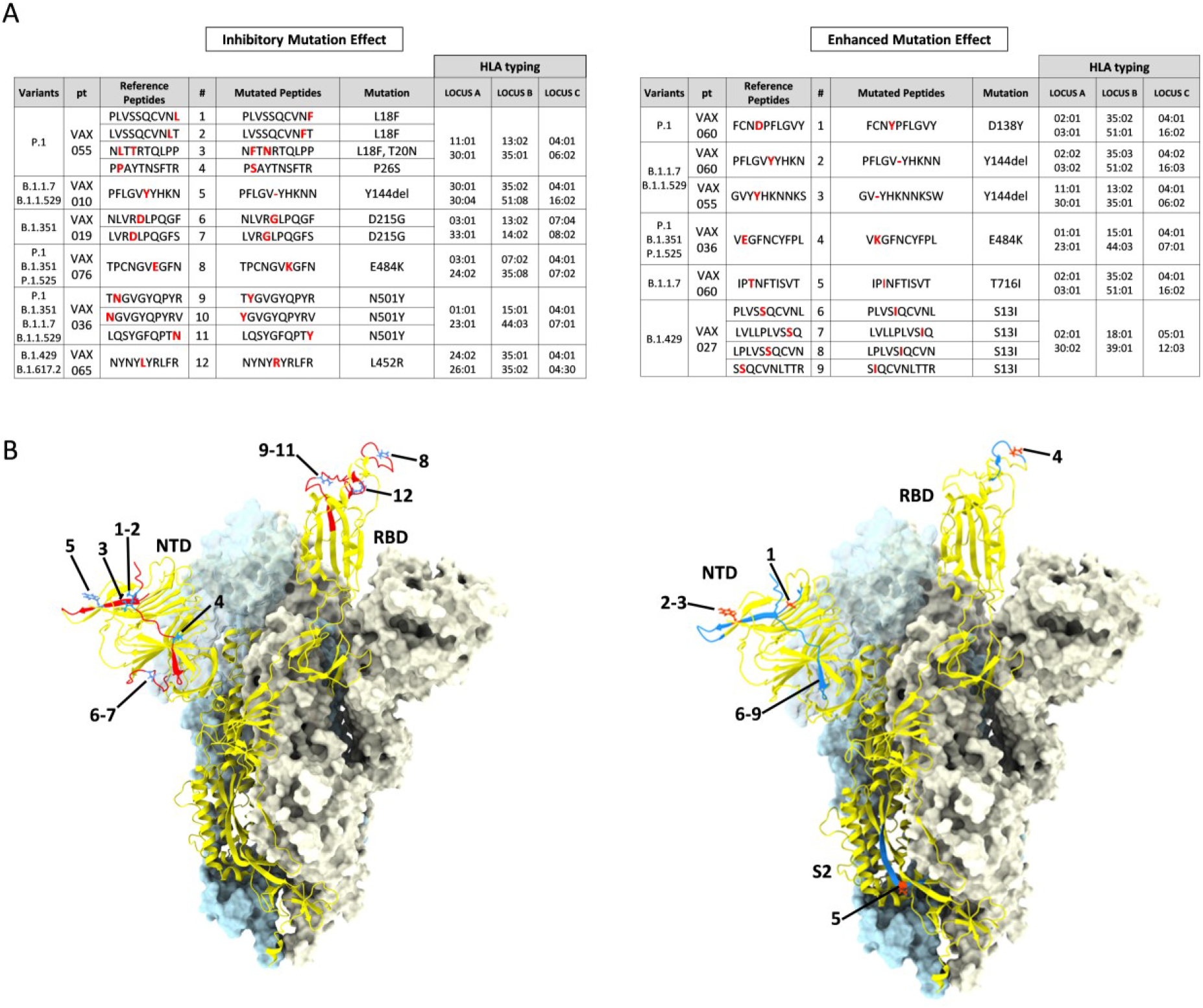
CD8 T cell variant epitopes and their location in the trimeric structure of the prototype spike protein ectodomain. **A)** List of mutated CD8 epitopes, that negatively (‘*Inhibitory’*) or positively (‘*Enhancing’*) modulate the CD8 T cell response compared to the corresponding prototype (‘Wuhan’) epitopes, identified by ex-vivo FluoroSPOT assay in vaccinated subjects (n=8); numbers in the # column allow to identify each peptide in the structure shown below. **B)** Trimeric structure of the prototype (Wuhan lineage) spike ectodomain (PDB 6ZXN) shown with one protomer (RBD-up) in *yellow cartoon* and the two other protomers (RBD-down) represented as *light blue* and *ice* surfaces. Inhibitory CD8 peptides (*left-side* panel) are shown on the *yellow* protomer as *red* ribbons, with the mutated residues displayed as *blue* sticks. The enhancing CD8 epitopes (*right-side* panel) are mapped on the *yellow* protomer as *blue* ribbons, with the mutated residues displayed as *red* sticks. Numbers (#) indicate the specific mutated peptides listed in panel (A); the substituted Ser13 residue of the enhancing peptides #6-9 is not displayed because the corresponding region in the spike protein structure is not resolved; NTD, N-terminal domain; RBD, receptor binding domain; S2, subunit 2.

### CD8 T cell reactivity against wild-type and mutated SARS-CoV-2 epitopes in convalescent subjects

To find out whether the immune potentiation effect of natural stimulatory mutations we observed in vaccinated subjects also occurs under natural SARS-CoV-2 infection, we tested PBMCs from 10 convalescent donors that were infected between April and October 2020, i.e., at a time when the variant viruses we selected for this study had not yet been detected in Italy. SARS-CoV-2 spike-specific CD8 T cell responses were detected 1-2 months after the acute phase of infection in 6 out of 10 convalescent subjects (Extended Data Table 4). Thirteen (45%) and nine (31%) of the positive peptide pools elicited inhibitory or enhanced responses, respectively (Fig. 5A), with a more prominent contribution of IFN-γ secreting T cells followed by double IFN-γ/TNF-α, IFN-γ/IL-2 and triple IFN-γ/TNF-α/IL-2 secreting T cells (data not shown). As in the case of vaccinated subjects, we were able to identify the optimal T cell epitope sequences within the peptide pools in all 3 convalescent subjects that we had the opportunity to test (Fig. 5B), which yielded concordant enhancing or inhibitory effects of individual mutations on T cell responses as assessed by FluoroSPOT or ICS assays both ex-vivo (Fig. 5C) and after in-vitro expansion (Fig. 5D). As indicated by the results of IFN-γ production and CD107a degranulation measurements (Fig. 5C, D), and in agreement with the analysis of T cell responses in vaccinees, immune-responses were sustained by CD8 T cells.

**Fig. 5.**
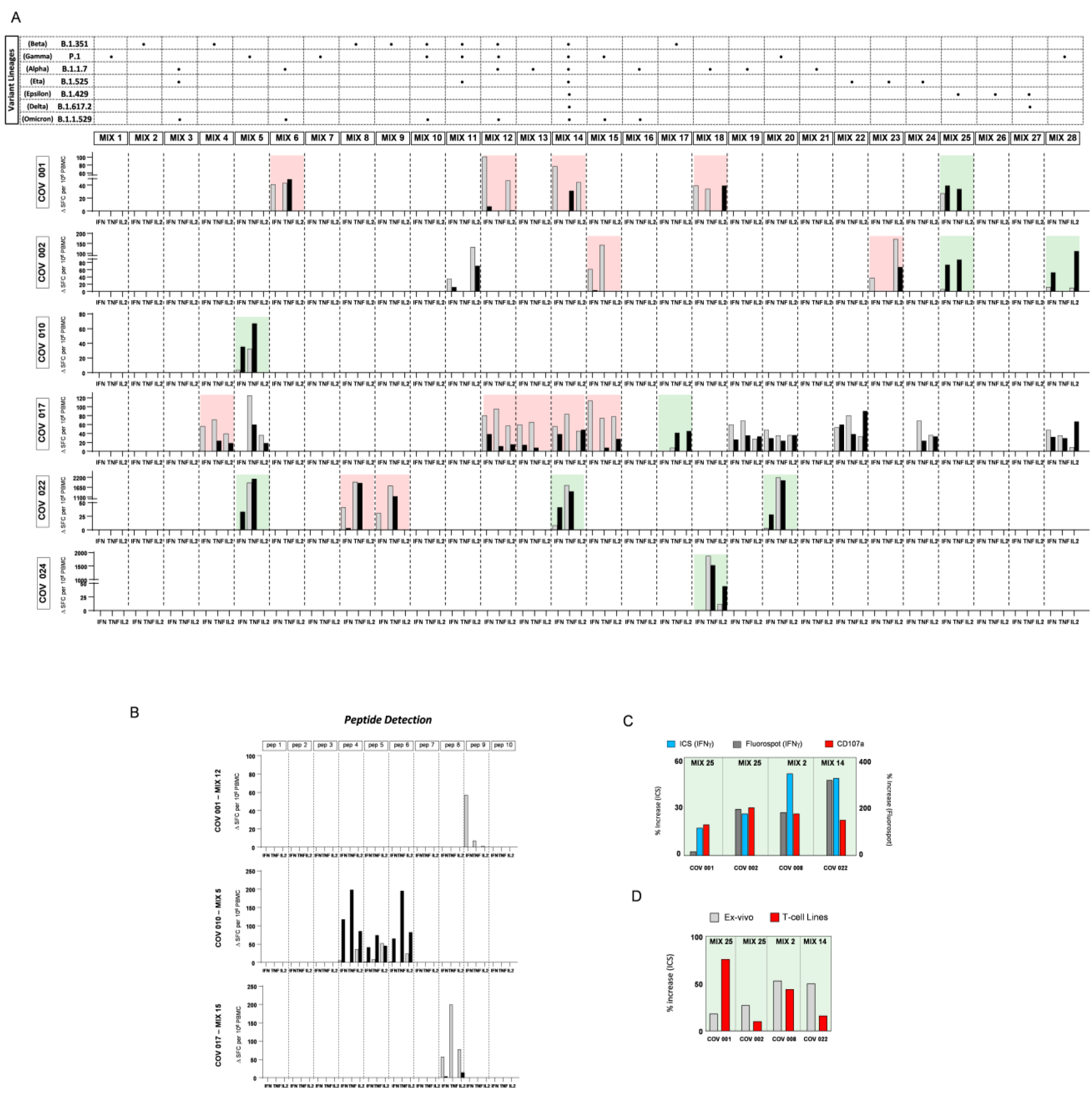
Effect of VOC and VOI mutations on spike-specific T cell responses in Covid-19 convalescent individuals. **A)** PBMCs from convalescent patients were stimulated for 18 hours by overlapping wild-type and variant peptides. Cytokine-secreting T cells were measured by FluoroSPOT assays. As in Fig. 1, only subjects with significant ex-vivo SARS-CoV-2 spike-specific CD8 T cell responses for at least two of the three analyzed cytokines (IFN-γ, TNF-α and IL2; 6 out of 10 tested patients) are illustrated (see Fig. 1 legend for further details). **B)** Individual stimulatory peptides identified by ex-vivo FluoroSPOT assays in three convalescent patients; *grey* and *black* bars represent the responses induced by wild-type and variant peptides, respectively. **C)** Bars represent the percentage increase of ex-vivo spike-specific IFN-γ production detected by FluoroSPOT (*grey bars*), ICS (*blue bars*) and CD8 T cell degranulation (*red bars*) upon PBMC stimulation with mutated *vs* wild-type spike peptide pools (n=4). **D)** Bars represent the percentage increase of IFN-γ detected by ICS either ex-vivo on PBMC (*grey bars*) or in vitro on expanded spike-specific CD8 T cell lines stimulated with variant *vs*. wild-type peptide pools (*red bars*) (n=4).

To gain initial insight on the possible mechanisms underlying the enhanced CD8 T cell reactivity properties, the nine peptides identified as ‘enhancing epitopes’ by our *in vitro* analysis were analyzed for HLA class-I binding, TCR recognition propensity and immunogenicity by two different prediction tools (IEDB and Prime 2.0; see ‘Methods’ for details). As shown in Extended Data Fig. 4, when compared with the corresponding prototype peptides, most of the experimentally validated enhancing 10-mer peptides were not recognized as highly immunogenic epitopes or high-affinity HLA/TCR binders, possibly because of their 10-mer length rather than the 9-mer length on which both prediction algorithms have been calibrated. In at least three cases (peptides #5, #6, #8; Extended Data Fig. 4), however, most or all prediction scores consistently indicated better HLA binding and TCR recognition properties for the mutated compared to the corresponding prototype peptides. There is also at least a case (#4) in which the mutation affects the P2 anchor thus determining a change in the HLA restriction element that co-occurs with a higher TCR functional avidity. The increased CD8 T cell reactivity of only a subset of the enhancing peptides may thus be explained by the random acquisition of superior HLA/TCR recognition properties.

### CD8 T cell responses to mutated SARS-CoV-2 epitopes in healthy uninfected/unvaccinated subjects

Next, we tested 11 unvaccinated and uninfected (‘naïve’) individuals to assess whether and to what extent cross-reactivity with pre-existing, SARS-CoV-2 unrelated memory T cells may contribute to anti-SARS-CoV-2 specific CD8 T cell responses. Such analysis may thus reveal whether cross-reactivity might explain the enhanced stimulatory effect exerted by some variant mutated peptides. Overall, 3% of the total stimulatory variant peptide pools induced detectable CD8 T cell responses in PBMCs from naïve subjects. More specifically, as shown in Extended Data Fig. 5, a significantly enhanced CD8 T cell response was observed against two different peptide pools in two of the 11 tested subjects.

A possible explanation for the CD8 T cell cross-reactivity observed in naïve subjects is previous exposure to peptides from other coronaviruses or unrelated human pathogens that share sequence homology with specific variant SARS-CoV-2 epitopes. We tested this hypothesis by using spike-associated SARS-CoV-2 T cell function enhancing epitopes as queries for a BLASTP search. Hits retrieved from this analysis were first restricted to 10-mer sequences at least 80% identical to specific SARS-CoV-2 peptides and further filtered to include only sequences from known human pathogenic microorganisms. None of the selected hits was found to be 100% identical to the mutated stimulatory SARS-CoV-2 peptides, whereas three mutated spike peptides (IPINFTISVT, LVLLPLVSIQ, PELGVYHKNN) shared 90% sequence identity with peptides from different bacterial, fungal and protozoan pathogens (see Extended Data Table 5). Several cases of 80% identity between additional function-enhancing spike variant peptides and SARS-CoV-2-unrelated human pathogens (including two infectious nematodes) also emerged from this analysis (Extended Data Table 5). Only one peptide (IPINFTISVT) was found to be similar to a subset of animal coronaviruses (not shown), while none of the common cold coronaviruses (CCCoV) reached the 80% identity threshold.

### Multispecificity of the CD8 T cell responses

Depending on the breadth and multispecificity of the whole repertoire of SARS-CoV-2-specific CD8 T cells, the overall protective antiviral CD8 activity may be affected to different extents by the loss or functional improvement of individual CD8-mediated responses caused by SARS-CoV-2 mutations. We addressed this issue by using two different peptide mixtures for PBMC stimulation: the first one composed of seven pools of 33 to 39 15-mer peptides overlapping by 10 residues and spanning the entire spike sequence; the second one comprising nine pools of peptides varying in length from 9 to 13 AA, previously described in the literature as CD8 T cell epitopes with different degrees of immunodominance and CD8 T cell stimulatory capacities (Extended Data Tables 6 and 7, respectively). A fraction of the latter CD8 epitopes is located within conserved Spike regions that are expected not to tolerate mutational changes because of structural/functional constraints. In addition, some of those epitopes are degenerate, i.e., able to associate with different HLA class I molecules and to be simultaneously presented to CD8 T cell populations of different HLA restriction^37^. Both mixtures were used in parallel to test the multispecificity of the response in all vaccinated and convalescent subjects that displayed decline or enhancement of spike-specific responses induced by variant mutations. In line with previous observations^38–40^, most vaccinated and convalescent donors exhibited multi-specific and powerful CD8-mediated T cell responses with simultaneous IFN-γ, TNF-α and IL-2 production stimulated by all seven 15-mer peptide pools (Fig. 6). Since each pool contains more than 30 peptides, this likely translates into dozens of CD4 and CD8 epitopes simultaneously recognized by T cells. In addition, most individuals also recognized a sizeable fraction of the immunodominant CD8 T cell epitopes previously identified within highly conserved spike regions. In contrast, a few subjects (n=4) recognized a more limited number of peptide pools (3 to 5) and displayed weaker responses to them. Collectively, these data indicate that in most, but probably not all subjects the presence of a vigorous and multi-specific CD8-mediated T cell response can largely compensate for the mutational loss of single responses.

**Fig. 6.**
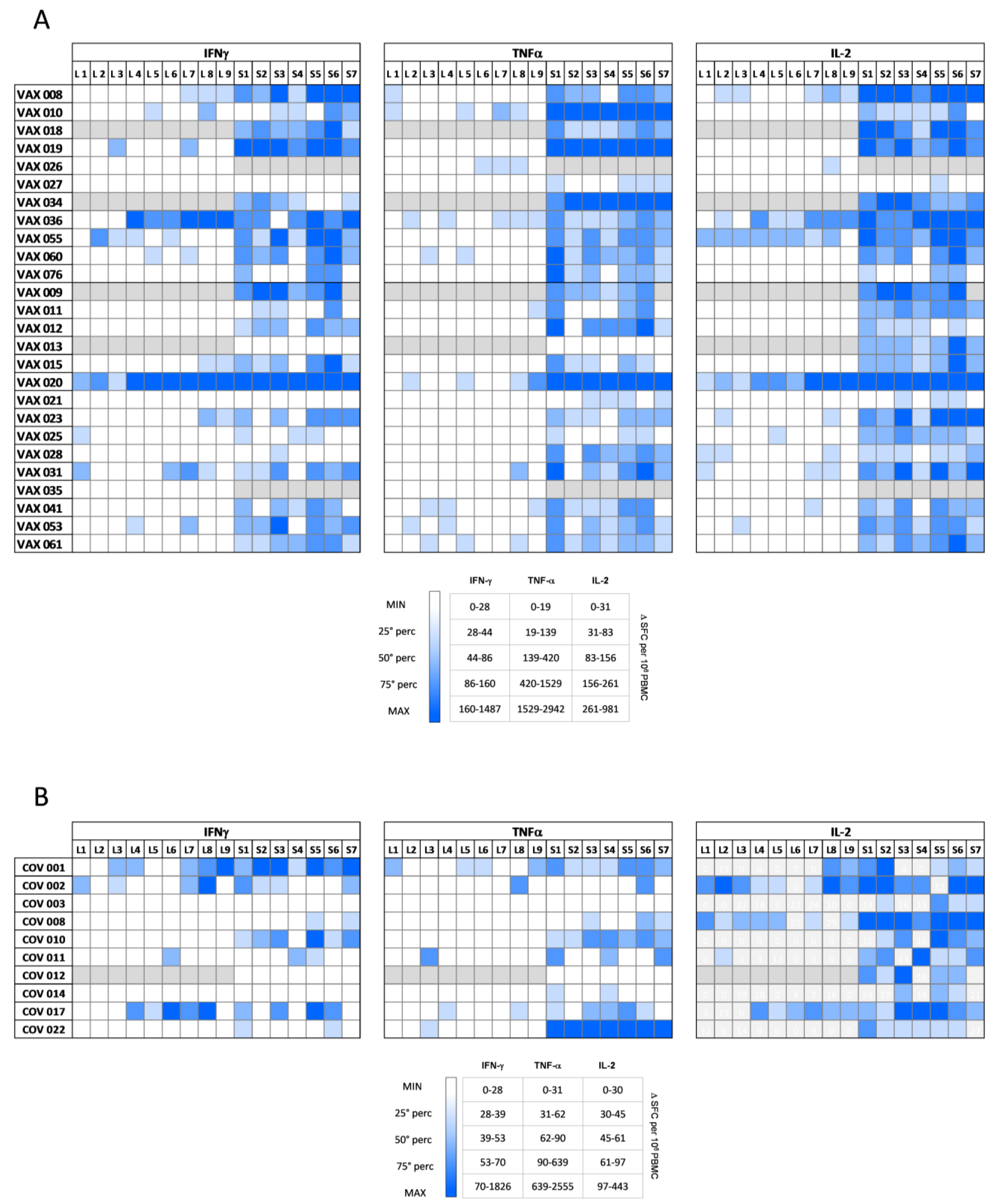
Breadth of the global spike-specific T cell response in vaccinated subjects and in SARS-CoV-2 infection convalescent patients. PBMCs from vaccinated (n=26) and convalescent (n=10) individuals were stimulated for 18 hours with seven pools of overlapping 15-mer peptides spanning the entire spike sequence (*right*) and nine pools of 10 peptides representing CD8 T cell epitopes previously described in the literature varying in length from 9 to 13 AA (*left*). Color intensity in the heatmaps indicates IFN-γ, TNF-α or IL-2 production levels measured as SFCs generated upon stimulation with each spike peptide pool (lower than 25^th^, 25^th^-50^th^, 50^th^ -75^th^, higher than 75^th^ percentile as specified in the inset) of individual vaccinated (**panel A**) and convalescent (**panel B**) subjects.

## DISCUSSION

Virus-neutralizing antibodies induced by prophylactic anti-SARS-CoV-2 vaccination are essential to prevent infection^18^, while elimination of virus-infected cells and cytokine-mediated purging of intracellularly hidden virus is the main role of CD8 T cells which is key to curb severe disease evolution^28,29,39,41^. Compared to short-lived antibody responses, CD8-mediated memory T cell immunity is long-lasting and remains detectable well after antibody waning^22,42^. Recognition of multiple epitopes per antigen and HLA allele, presentation of individual peptides by multiple HLA molecules and location of target epitopes in structurally/functionally constrained regions of SARS-CoV-2 antigens represent CD8 T cell features which are crucial for counteracting SARS-CoV-2 immune evasion.

Several studies have analyzed the effect of SARS-CoV-2 mutations on CD8 T cell-mediated protection by comparing the immune responses elicited by mega-pools of synthetic peptides derived from the parental Wuhan SARS-CoV-2 strain and from the several VOCs/VOIs viral lineages that have emerged since the beginning of the COVID-19 pandemic ^22–25^. At variance with most previous studies, here, we analyzed the effect of a large set of spike-associated (VOC/VOI) mutations on CD8 T cell activity in relation to the breadth and quality of the overall CD8 responses in vaccinated and convalescent individuals. Our assumption was that individual mutations should really affect anti-viral CD8-mediated protection only in the context of an extremely narrow repertoire of CD8 specificities.

To maximize the likelihood of identifying functionally relevant mutated epitopes, we designed multiple panels of synthetic peptides of optimal length for CD8 T cell recognition spanning the prototype and the corresponding VOC and VOI regions containing the most relevant mutations that have accumulated overtime. CD8 T cell responses were further characterized by two additional peptide panels, one comprised of 96 validated CD8 epitopes of varying HLA restriction representing the majority of the CD8 epitopes within the spike region so far described in the literature and the other composed of 15-mer peptides overlapping by 10 residues covering the entire spike sequence. All these pools were tested in parallel in individual vaccinated and convalescent subjects.

All variant peptide pools were recognized by at least one of the 17 vaccinated responders (out of 36 tested subjects), with some vaccinees reactive to multiple mutated peptide pools. As expected, most SARS-CoV-2 mutations (~70%) were either inhibitory or neutral, meaning that the corresponding peptides elicited CD8 immune responses lower than, or comparable to, those induced by the corresponding prototype epitopes. Interestingly, however, ~30% of the variant epitopes outperformed the CD8 reactivity of the corresponding prototype epitopes or elicited immune responses that were otherwise undetectable with prototype non-mutated epitopes, thus behaving as naturally evolved heteroclitic peptides.

Heteroclitic peptides are typically created artificially, to induce stronger T cell responses by changing specific HLA anchor residues in order to increase HLA binding affinity or to generate new HLA anchor motifs, but also by modifying TCR binding residues in order to increase the affinity of peptide-HLA recognition by TCRs^43–46^. SARS-CoV-2 mutations within epitopic or flanking sequences able to mediate a more robust stimulation of CD8 T cell responses than their wild-type counterparts have occasionally been reported but never investigated in detail^47–50^. Of note, we detected function-enhancing amino acid changes in all tested variants and some of the mutated peptides induced opposite (stimulatory or inhibitory) effects in different vaccinated subjects (e.g., the Y144 deletion was inhibitory in VAX 010 but enhancing in VAX 060), an inter-subject variability that is likely explained by the marked individual TCR plasticity and by a possible differential effect of mutations on HLA binding if the same peptide can be presented to CD8 T cells by different HLA molecules in different subjects.

Multiple lines of evidence in our study indicate that the effects of mutated variant epitopes, especially the heteroclitic-like ones, represent a real modulation of T cell functions of potential relevance to anti-viral protection, rather than just an in vitro phenomenon. First, the effects of individual mutations were classified as stimulatory or inhibitory based on a stringent effect size analysis, using the large effect threshold as a discriminant to designate T cell responses falling below or above this cutoff as decreased or increased significantly. Moreover, only responses with a concordant modulation of at least two of the three tested cytokines were considered as reliable indicators of a positive or negative effect. Second, the same stimulatory or inhibitory effects detected in the initial screening with peptide pools were reproduced by individual immunogenic peptides identified within each stimulatory peptide pool and were confirmed by dose titration experiments comparing individual variant with the corresponding wild-type peptides. Third, mutated peptide effects initially detected by multi-cytokine FluoroSPOT assays were independently validated by ex-vivo intracellular cytokine staining and CD107 degranulation, which confirmed that the observed responses were indeed CD8 T cell-mediated and that the modulatory effect also affected the cytolytic component of the CD8 function. The fact that these CTL responses are entirely sustained by CD8 T cells with no detectable contribution by CD4 T cells, was further corroborated by the concordant results obtained with in-vitro expanded T cells stimulated with mutated or prototype peptides. Importantly, similar stimulatory and inhibitory effects were observed in 6 out of 10 convalescent patients, indicating that positive and negative modulation of CD8 responses also occurs during natural infection and may be clinically relevant.

Decline or abrogation of CD8-mediated T cell responses by newly emerged, CD8 epitope-targeting mutations is an expected and well documented event for infections caused by highly variable RNA viruses, which may lead to mutated viral escape if the breath of the overall antiviral CD8-mediated response is sufficiently narrow to preclude the compensatory role of co-existing non-mutated immunodominant epitopes. Since CTL responses to SARS-CoV-2 have generally been reported to be broadly multi-specific^38–40^, it is usually thought that virus escape from CD8 T cell control is a rare event requiring the concomitant emergence of multiple inactivating mutations targeting different immunodominant epitopes to make the virus invisible to all or most protective CTL clones. Our data, however, indicate that in a fraction of vaccinated and SARS-CoV-2 infected subjects (e.g., VAX 026, VAX 027, VAX 013, VAX 021, COV 003, COV 014) CD8 T cell responses appear to be focused on a limited number of spike regions, suggesting the possibility that in this more restricted setting even the abrogation of single CTL responses may negatively impact the overall CTL-mediated anti-viral protection. It is also important to note, however, that in natural infection CTL responses are stimulated by other SARS-CoV-2 antigens in addition to the spike protein, thus making CD8 T cell multi-specificity even wider and further reducing the possibility of a mutational effect on the overall CD8 T cell-mediated antiviral protection^38,40,51–53^.

Conversely, it is less clear and counterintuitive the appearance of CD8 T cell function-enhancing (“suicide”) mutations, that may potentiate protective immunity by facilitating virus elimination rather than its persistence. Since CD8 response-enhancing peptides were also identified in a fraction (2 out of 11) of naïve subjects, suggesting the existence of some cross-reactivity with pre-existing memory CD8 T cells of different specificity, we asked whether the enhancing effect of these mutated peptides might reflect sequence homology with polypeptides from unrelated human pathogens. Interestingly, multiple 10-mer peptides, 80% to 90% identical to spike stimulatory peptides, were identified within polypeptide sequences from unrelated human pathogens contained in the NCBI repository. This points to the potential relevance of SARS-CoV-2-specific CD8 T cell cross-reactivity triggered by pre-existing memory responses raised against unrelated microorganisms in shaping the strength and quality SARS-CoV-2-specific CTL responses^54,55^. These homology data are also in line with the frequent detection of SARS-CoV-2-specific T cells in virus-unexposed individuals^38,40,56–60^. This potential heterologous priming, however, is not unique to the enhancing peptides, as comparable numbers of unrelated homologous sequences sharing similar levels of amino acid identity with SARS-CoV-2 epitopes were retrieved by a BLASTP search performed using the inhibitory peptides as queries (data not shown). Furthermore, the enhancing effect per se will necessarily result in a better CD8 T cell reactivity, which would thwart variant virus selection.

An alternative hypothesis is that selection of such variant enhancing epitopes is primarily determined by mutations that while causing specific structural/functional alterations ultimately favoring virus transmissibility, fortuitously hit spike sequences of potential relevance for CD8 T cell activation. This is supported by two lines of evidence: first, our epitope mapping data with overlapping peptides indicates that all selected variant mutations are contained within CD8 T cell epitopes; second, variant epitope mapping on the three-dimensional structure of the spike protein indicates that nearly all mutated epitopes are located within functionally crucial spike regions, such as the receptor binding and the N-terminal domains of the spike S1 subunit, involved in ACE2 receptor engagement and in co-receptor recognition, which can have profoundly favorable effects on virus fitness. This is in line with previous reports in other virus infections indicating that individual amino acid changes can exert multiple and seemingly divergent effects, with a possible concomitant impact on both antigenicity and cell tropism^61^. In this scenario, the positive effect of SARS-CoV-2 mutations on virus fitness would allow the virus to gain a selective advantage of general relevance for the overall human population and would thus represent the primary driver of variant virus selection and spreading, regardless of the specific host genetic background and individual CD8 T cell reactivity. Only upon infection of individuals carrying the appropriate HLA restriction element for peptide presentation, CD8 T cells would be activated and mutations would affect their reactivity. The random positioning of the amino acid changes within the epitope sequence and the local chemical modifications imposed by the variant amino acid would thus determine the positive or negative impact on HLA and/or TCR binding. In keeping with this possibility, we found that a fraction (3 out of 9) of the mutated enhancing peptides displays predictive scores for HLA/TCR interaction higher than those of the corresponding prototype peptides (Extended Data Fig. 4).

These enhancing mutations might be irrelevant to the overall anti-viral protection in the presence of widely multispecific CD8 T cell responses, but might bear pathogenetic relevance if the responses are more narrowly focused, as we observed in some vaccinated subjects. Thus, individual SARS-CoV-2 variant mutations can exert opposite effects on the spread and progression of infection: an increase in CD8 T cell activity making SARS-CoV-2 more vulnerable to cytotoxic responses would be the price to be paid in a limited proportion of patients for the virus to acquire a greater general advantage at the population level resulting from an improved cell tropism and/or transmissibility.

## METHODS

### Study participants

36 health care workers (HCW) were enrolled into a vaccination study at the Unit of Infectious Diseases and Hepatology of the Parma University Hospital, Italy. PBMCs and serum samples from participants were collected 1-2 months after the second dose of Pfizer/BioNTech BNT162b2 vaccination. At the same time point of sample collection, Nucleocapsid (N) immunoglobulin G (IgG) was tested by CMIA (chemiolumescent microparticle immunoassay; Architect-Abbot) to exclude previous SARS-Cov-2 infection (1 positive donor out of 36). 12 patients were also recruited from April to October 2020 after hospitalization for SARS-CoV-2 infection, as indicated by positive RT-PCR. Samples from convalescents were obtained 1-2 months after disease onset. The study was approved by the competent local Ethic Committee and all patients provided written, informed consent. Details on the vaccinated and convalescent donors are reported in Extended Data Tables 1 and 4.

### Sequence analysis and peptide design

Genome sequences derived from the different VoCs and VoIs were compared with the original Wuhan SARS-CoV-2 strain by the GISAID data base in order to map all spike amino acid substitutions and deletions associated with the main Variants of Concern and Variants of Interest, including the UK (Alpha), South-African (Beta), Brazilian (Gamma), Californian (Epsilon), Indian (Delta) VoCs, as well as Nigerian (Eta) VoIs (GISAID data base - https://cov.lanl.gov/content/index) (Extended Data Table 2). Next, we designed a set of peptides of optimal length for cytotoxic T cell recognition, i.e. 10 AA long, overlapping by nine residues, with variable numbers of flanking AA at both sides of the mutations. For each relevant mutation we synthesized 2 pools of 10 peptides (a wild type and a mutant pool, except for mix 1 consisting of 18 peptides), containing individual mutations at each position of the peptides, for a total of 28 couples of pools (Extended Data Tables 2 and 3). Our study was already in progress when the delta and omicron variants emerged; thus, most delta and omicron mutations were not included, with the exception of those shared with other variants.

To investigate the breadth of the spike-specific T cell response elicited by vaccination, PBMCs were also stimulated with 15-mer peptides overlapping by 10 amino acids spanning the overall Wuhan spike sequence and consisting of 7 pools, each composed of about 40 peptides (Extended Data Table 6).

Finally, to characterize further strength and multispecificity of spike-specific CD8 T cell responses primed by vaccination, we used an additional set of 96 peptides, previously described in the literature as Spike-specific CD8 T cell epitopes, varying in length from 9 to 13 AA (Extended Data Table 7). A proportion of these peptides are located within conserved spike regions, as defined by Shannon entropy (SE), which is a measure of variability of genetic mutations^37^ for each amino acid position, calculated on a multiple alignment of spike protein sequences. Decreasing entropy thresholds values of 0.0025, 0.001 and 0.0005 were defined by preliminary analysis of 1,400,000 million spike protein sequences retrieved (as of June 1st 2021) from the daily updated GISAID database^37^. Three levels of amino acid (AA) residue conservation were then arbitrarily defined, allowing us to distinguish ‘conserved’, ‘highly conserved’ and ‘hyper-conserved’ spike regions (Extended Data Table 7) and to map each epitope to variably constrained portions of the spike protein. Based on this analysis 25 peptides are located within conserved, 8 within highly conserved and 4 within hyper-conserved Spike regions.

### HLA-typing analysis

High resolution HLA typing of all samples was performed using different NGS kits. All samples were genotyped for 11 HLA loci, namely HLA-A, -B, -C, -DRB1, -DRB3, -DRB4, and -DRB5, -DQA1, -DQB1, -DPA1, -DPB1. Pooled libraries were loaded on a iSeq 100 Reagent v2 (cartridge and flow cell, Illumina). Paired-end sequencing was performed on the iSeq100 next-generation sequencing (NGS) platform (Illumina), 151 cycles in each direction. HLA typing analyses were made using different HLA typing software packages (AlloSeqTM Assign®, CareDx; TypeStream Visual, One Lambda; NGSengine®, GenDx) along with the current version of the IPD-IMGT/ HLA database.

### SARS-CoV-2 pseudoviruses generation and neutralization assay against the original viral strain and variants

Lentiviral vector-based SARS-CoV-2 S pseudovirus was generated as previously described with minor modifications^62^. Sub confluent HEK 293T cells were cotransfected with pLV-EF1α-(turboGFP-Luc2)-WPRE transfer vector, p8.74 packaging vector, pseudotyping vector coding for Spike glycoproteins (Wuhan-Hu-1; B.1 Lineage, China) and pREV with PEI (Polysciences, Inc., Warrington, PA, USA) (1 mg/mL in PBS) (ratio 1:2.5 DNA/PEI). Transfected cells were incubated for 48 h at 37 °C and 5% CO2. The flask was then frozen–thawed at −80 °C; transfected cell supernatant (TCS) containing S pseudovirus was clarified via centrifugation, filtered through a 0.45 μm filter (Millipore, Merk, Darmstadt, Germany), aliquoted, tittered by limited dilution and stored at −80 °C. Serum neutralization assay was performed as previously described^62^. S pseudovirus preparation diluted in complete EMEM with 10% FBS was added to each well containing the diluted sera and left to incubate at room temperature for 1.5 h. Next, 104 HEK/ACE2/TMPRRS2/Puro cells, were added to each well and left for 60 h at 37 °C and 5% CO2. Luciferin was added to each well just before the reading of the microplate with the luminometer (Victor, Perkin Elmer). Neutralization titer 50 (NT50/ml) was expressed as the maximal dilution of the sera where the reduction of the signal is ≥50%. Each serum was tested in triplicate.

### Peripheral blood mononuclear cell isolation

Peripheral blood mononuclear cells (PBMC) were isolated from fresh heparinized blood by Ficoll-Hypaque density gradient centrifugation and cryopreserved in liquid nitrogen until the day of analysis.

### FluoroSPOT assay

IFN-γ/TNF-α/IL-2 three-colour FluoroSPOT assay was performed using a panel of peptides (10-mers overlapping by 9 aa) pooled in 56 mixtures containing most of the VOC and VOI mutations within the spike region (Extended Data Table 3). The day before the assay, the plates were activated by adding 70% ethanol followed by overnight incubation with IFN-γ/TNF-α/IL-2 capture antibodies. After decanting the plate, 2-4 × 10^5^ PBMCs per well were seeded in duplicate in *CTL Medium* and Spike-specific T-cell responses were analyzed after overnight incubation with individual peptide mixtures (1μM) as IFN-γ/TNF-α/IL-2 production according to the manufacturer’s instruction (CTL, Europe, Germany). Spots were counted using an automated reader system (ImmunoSpot Ultimate UV Image analyzer, CTL Europe, Germany). Cytokine-secreting cells were expressed as spot forming cells (SFC) per 1 × 10^6^ cells after subtraction of the background. Positive controls consisted of PBMCs stimulated with CMV, EBV and influenza peptides. FluoroSPOT was considered positive if the number of spots in the stimulated wells was at least 3 standard deviations above background and the difference between the number of spots in the stimulated and unstimulated wells was above 10.

### Flow cytometry analysis

PBMC or expanded T cell lines were stimulated for eighteen hours at 37°C with or without SARS-CoV-2 peptide pools (1μM) as mentioned above in the presence of Brefeldin A (GolgiPlug 1 μL/mL) and Monensin (GolgiStop 0.5 μL/mL). Cells were washed and surface-stained with Live/Dead fixable dead cell stain kit (Invitrogen) and anti-CD3, anti-CD4, anti-CD8 (all from BD Biosciences) fluorochrome-conjugated antibodies. PBMC were then fixed, permeabilized using the Fix and Perm kit (Nordic MUbio) and stained with anti-IFN-γ (BD Biosciences) and anti-TNF-α (Miltenyi) conjugated mAbs for the detection of intracellular cytokines, and using an anti-CD107a antibody (BD Biosciences) for the study of the cytotoxic potential. Samples were acquired on a BD LSR Fortessa and analyzed with the FlowJo software (BD Bioscences).

### Expanded T cell lines

PBMC were stimulated with selected SARS-CoV-2 peptide pools (1μM) at 37°C in RPMI medium supplemented with 8% human serum and cultured for 7-9 days in the presence of 50 UI/ml of recombinant IL-2 (R&D System).

### Variant peptide mapping on the three-dimensional structure of the spike protein

Variant SARS-CoV-2 peptides identified in vaccinated individuals by ex-vivo FluoroSPOT analysis were mapped on a high-resolution, one RBD-up spike structure (PDB: 6ZXN)^63^ using the ChimeraX software^64^.

### Homology analysis of variant SARS-CoV-2 peptide with unrelated human pathogen sequences

The search for protein sequences of other organisms containing fragments identical or similar to the SARS-CoV-2 variant enhancing peptides was performed as follows. For every peptide used as a query, Blastp analysis (https://blast.ncbi.nlm.nih.gov/Blast.cgi?PAGE=Proteins) within the non-redundant protein sequences (nr) database, excluding the organism “Severe acute respiratory syndrome coronavirus 2 (taxid:2697049)” and choosing PAM30 matrix plus other default parameters, allowed the isolation of 1000 sequences. Among them, a selection was done using the following criteria: i) sequences still correlated with human respiratory syndrome coronavirus 2 or other animal coronavirus were eliminated; ii) after a multiple alignment of all identified peptides with the corresponding query, using ClustalW program, the peptides with an inserted gap bigger than one amino acid were excluded, as well as all peptides with less than 8 amino acid residues identity; iii) identical peptides derived from the same protein sequences belonging to the same family of micro-organisms were grouped when repeatedly reported in order to consider only a single peptide. The remaining protein sequences were further subjected to selection in order to consider only those corresponding to microorganisms that may have come into contact with, or infected, humans and therefore be capable to trigger an immune response.

### Statistical analysis and HLA affinity/MHC binding/immunogenicity prediction tools

Data were analyzed by GraphPad Prism (GraphPad Software, La Jolla, CA). Statistical significance was assessed by the Mann-Whitney U test for non-paired samples and the Wilcoxon signed rank test for paired data. Differences between multiple patient groups were evaluated by the non-parametric one-way ANOVA test ccrrected for pairwise multiple comparisons. Frequencies were compared by Chi Square with Yates’ correction.

To establish relevant differences between T cell responses stimulated by wild type and mutated peptides we applied an *Effect size* calculation. Fold changes can be assimilated to Risk Ratios (RR) for which their use as “effect size” measures has been suggested in the literature^65^. Effect size reveals how relevant is the relationship between variables or the difference between groups. Measures of effect size are usually divided into “small effect", “medium effect” and “large effect”. A large effect size means that a research finding has practical significance, while a small effect size indicates limited practical implications. For our data we chose the “large effect” threshold (RR = 3.0), as the discriminant for the most important values. Taking the logarithm of this value, we defined the corresponding thresholds in the log scale (0.48). Specifically, for each T cell response stimulated by a mutant pool, we calculated the fold-change relative to the ancestral pool (mutated/wild-type). Only T cell responses below or above the “*Large Effect size”* values were considered significantly decreased or increased (< −0.48 or > +0.48).

PRIME 2.0 and IEDB prediction tools were used to predict HLA affinity, MHC binding and immunogenicity of wild type and mutated spike peptides positively modulating CD8 T responses. The PRIME 2.0 immunogenicity predictor tool combines peptide TCR recognition propensity with predicted affinity to HLA-class-I molecules. The lower the percentile rank score PRIME value, the higher the peptide expected ability to elicit CD8 T cell recognition. Prime % score values lower than 0.5% threshold represent good HLA-I binding candidates^66^. The IEDB Class I Immunogenicity tool (https://www.iedb.org/) was employed to predict the immunogenicity score and the binding affinity (IC_50_) of peptides for HLA class I molecules. A higher immunogenicity score indicates a greater probability of eliciting an immune response. The HLA-I binding tool predicted output is IC_50_nM.

## ACKNOWLEDGEMENTS

We thank all of the patients and control volunteers who participated in this study and all of the clinical staff who helped with recruitment and sample collection; we thank Rosa Sorrentino (Department of Biology and Biotechnology Charles Darwin, Sapienza University, Rome, Italy) for critical reading of the manuscript and helpful discussion.

## AUTHOR CONTRIBUTIONS

C.T., A.V., M.R.: execution of experiments, acquisition of data, statistical analysis, analysis and interpretation of data. D.C., A.B.: contribution to figure drawing, analysis and interpretation of data and contribution to the selection of the references to quote. D.L.: administrative support. L.S., F.B., T.M., A.T., A.N.: recruitment and characterization of the patients. G.D.: neutralization assay. P.Z., M.B., S.G.: HLA typing analysis. P.F., I.M., S.U.: execution of experiments and interpretation of data. G.P.: statistical analysis. G.M., A.M., D.C., S.O.: critical revision of the manuscript. C.F., C.B.: study concept and design, critical revision and editing of the manuscript, obtained funding, study supervision and interpretation of data. All authors contributed to the article and approved the submitted version.

