## Extended figure and table for "Natural heteroclitic-like peptides are generated by SARS-CoV-2 mutations"

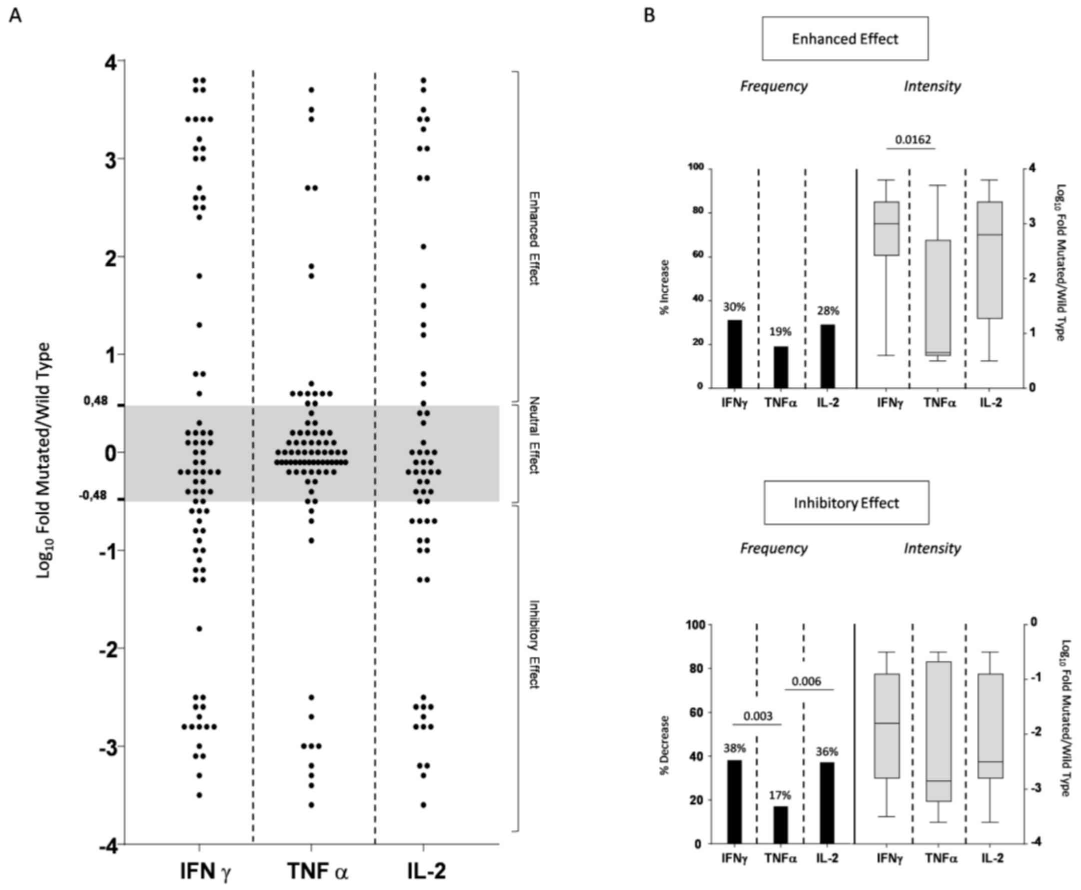


**Extended Data Fig. 1. Effect of SARS-CoV-2 variant-associated mutations on spike-specific T cell responses of vaccinated donors. A)** Fold-change difference between ex-vivo IFN-γ-, TNF-α- and IL-2- secreting T cell responses induced by mutated and wild-type (Wuhan type) peptides detected by a FluoroSPOT assay in vaccinated individuals (n=17). All FluoroSPOT responses shown in Fig. 1 are represented. The area delimited by a *grey* background (comprised between +0.48 and -0.48; |0.48| = large-effect size threshold) corresponds to neutral CD8 T cell responses (i.e., SFC values induced by variant mutated and WT peptide stimulation are not significantly different), while T cell responses below or above the *grey* area represent significantly decreased or increased CD8 T cell responses elicited by variant peptides. Each dot corresponds to the fold change response induced by each mutated compared to the corresponding wild type peptide. **B)** Frequency and intensity of the enhancing (*top*) and the inhibitory (*bottom*) effects of variant peptide pools on IFN-γ, TNF-α and IL-2 responses. Frequency values (*left part* of each panel) were evaluated statistically using a Chi-square test with Yates' correction. The intensity of the enhancing and inhibitory effects (*right part* of each panel) is expressed as median fold-change values (statistically evaluated by one-way ANOVA with Dunn's multiple comparisons test) of T cell responses stimulated by variant and wild-type peptides.

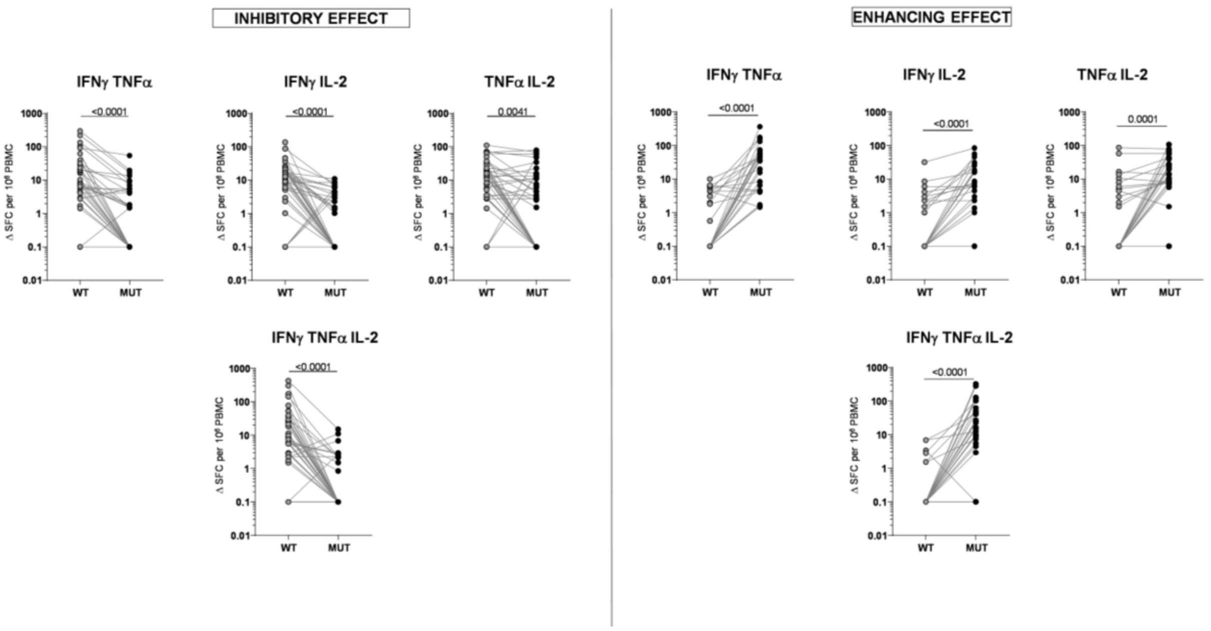


**Extended Data Fig. 2. Polyfunctionality of spike-reactive CD8 T cells.** Frequencies of T cells capable of simultaneously producing multiple cytokines upon Wuhan prototype (WT) or variant spike peptide pool stimulation in vaccinated responders (n=17); data representing diminished or enhanced T cell responses are grouped in the *left* (‘*inhibitory effect’*) and in the *right* (‘*enhancing effect’*) panels. Each circle (*grey* or *black*, for wild-type or variant peptide pools, respectively) represents the frequency of double and triple positive cytokine-secreting CD8 T cells; statistical analysis was performed by the Wilcoxon matched-pairs signed rank test.

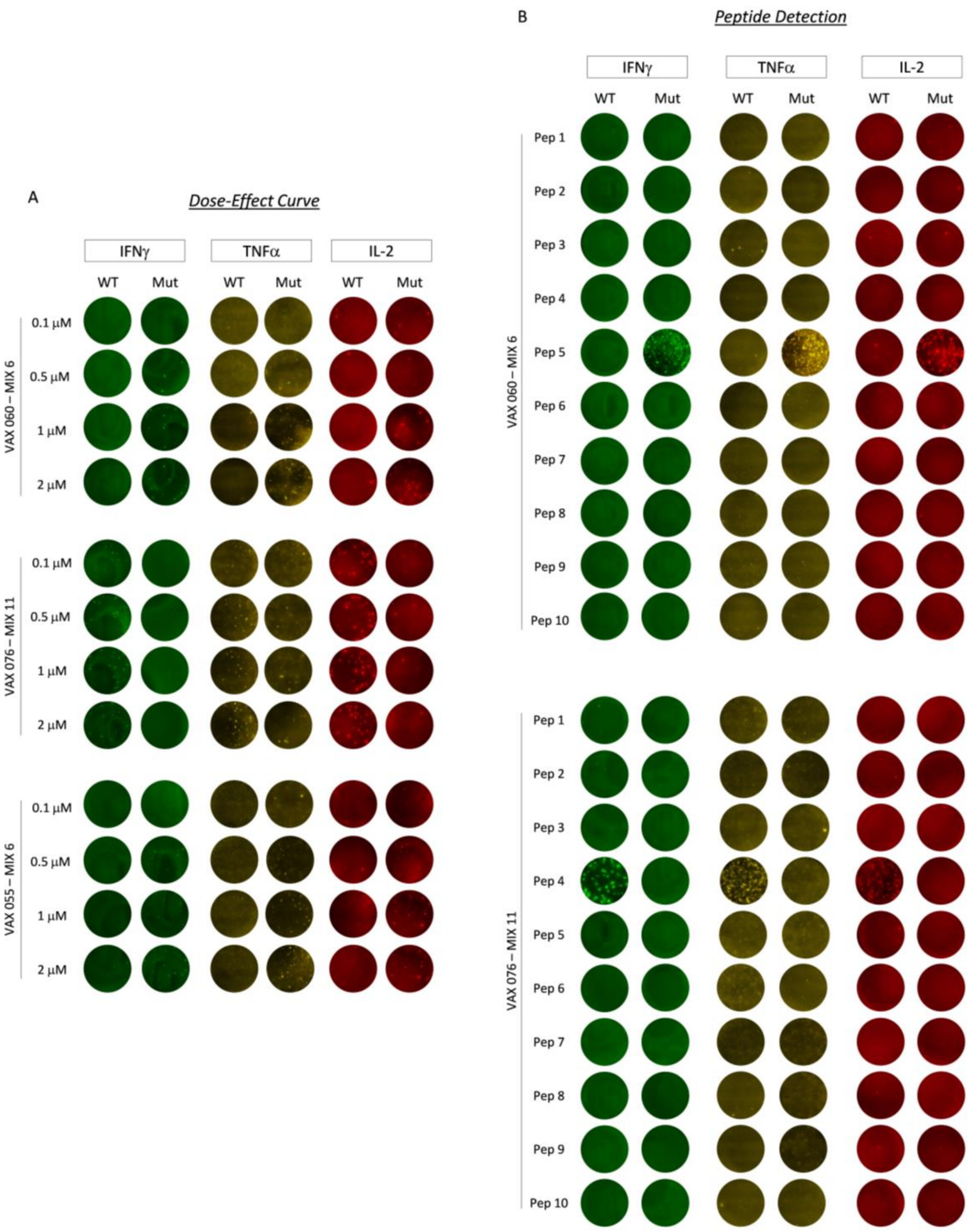


**Extended Data Fig. 3. Representative FluoroSPOT assay data. A)** Examples of dose-response curves determined by ex-vivo FluoroSPOT assays performed on PBMCs from three representative vaccinated donors stimulated with different peptide concentrations. **B)** Representative ex-vivo FluoroSPOT assays for the identification of specific CD8 T cell response-enhancing (top) or inhibitory (bottom) peptides contained within the peptide pools.

| | | HLA | | | IEDB tools | | | | | | | Prime 2.0 | | |
|---|---|---|---|---|---|---|---|---|---|---|---|---|---|---|
| Variants | Vaccinee | LOCUS A | LOCUS B | LOCUS C | # | Prototype peptides | Prot. peptide IC50 class (HLA) | Mutated peptides | Mut. peptide IC50 class (HLA) | Mutated Peptide IC50 variation | Immunogenicity score variation (Δ) | Prot. peptide ranking (%) (HLA/TCR) | Mut. Peptide ranking (%) (HLA/TCR) | Mutated peptide score variation (%) |
| P.1 | VAX 060 | 02:01<br>03:01 | 35:02<br>51:01 | 04:01<br>16:02 | 1 | FCNDPFLGVY | 35:02 | FCNYPFLGVY | 35:02 | ↑ | -0.03 | 16:02 | 51:01 | |
| B.1.1.7<br>B.1.1.529 | VAX 060 | 02:02<br>03:02 | 35:03<br>51:02 | 04:02<br>16:03 | 2 | PFLGVYYHKN | 03:02 | PFLGV-YHKNN | 02:02 | ↑ | -0.175 | 02:02 | 02:02 | |
| | VAX 055 | 11:01<br>30:01 | 13:02<br>35:01 | 04:01<br>06:02 | 3 | GVYYHKNNKS | 30:01 | GV-YHKNNKSW | 30:01 | ↑ | -0.178 | 11:01 | 11:01 | |
| P.1<br>B.1.351<br>P.1.525 | VAX 036 | 01:01<br>23:01 | 15:01<br>44:03 | 04:01<br>07:01 | 4 | VEGFNCYFPL | 44:03 | VKGFNCYFPL | 23:01 | ↑ | 0 | 44:03 | 07:01 | |
| B.1.1.7 | VAX 060 | 02:01<br>03:01 | 35:02<br>51:01 | 04:01<br>16:02 | 5 | IPTNFTISVT | 35:02 | IPINFTISVT | 35:02 | ↓ | 0.03 | 35:02 | 51:01 | |
| B.1.429 | VAX 027 | 02:01<br>30:02 | 18:01<br>39:01 | 05:01<br>12:03 | 6 | PLVSSQCVNL | 02:01 | PLVSIQCVNL | 02:01 | ↓ | 0.29 | 02:01 | 02:01 | |
| | | | | | 7 | LVLLPLVSSQ | 02:01 | LVLLPLVSIQ | 02:01 | ↓ | 0 | 30:02 | 30:02 | |
| | | | | | 8 | LPLVSSQCVN | 30:02 | LPLVSIQCVN | 18:01 | ↓ | 0.29 | 18:01 | 18:01 | |
| | | | | | 9 | SSQCVNLTTR | 30:02 | SIQCVNLTTR | 30:02 | ↑ | 0 | 30:02 | 02:01 | |

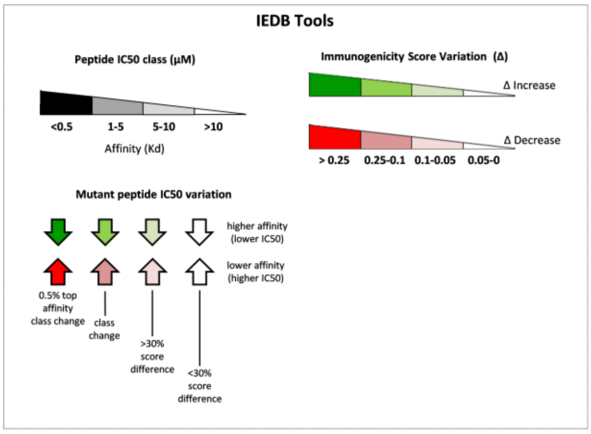


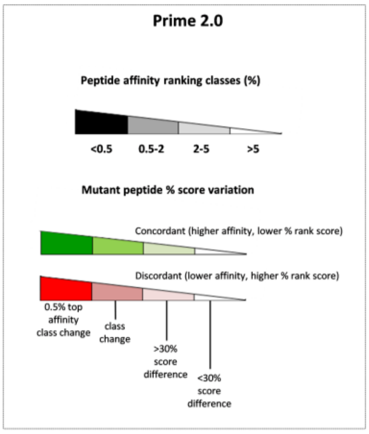

**Extended Data Fig. 4. HLA binding, TCR recognition and CD8 T-cell immunogenicity parameters of the enhancing peptides evaluated with two different prediction tools.** IC50 values predicted with the IEDB tools are shown on a background of decreasing grey color intensities, which, as explained in the scale caption at the bottom, correspond to decreasing HLA affinities (given as dissociation constants, Kd); IC50 prediction was based on the specific HLA alleles of individual vaccinees (only the values for the best-scoring alleles are reported). The direction of the arrows in the 'Mutated Peptide IC50 variation' column indicates a higher (upward) or a lower (downward) IC50 value of the mutated peptides with respect to the corresponding prototype peptides. The range of variation is indicated by the different shading intensities of the green (higher affinity) or red (lower affinity) colors of the arrows, as specified in the scale caption. Variations of the immunogenicity scores are indicated by the green (increase relative to the corresponding prototype peptide) or red (decrease) color of the boxes in which the variation (Δ) values are displayed; color intensity is proportional to the extent of variation as specified in the scale caption.

The last three columns summarize the results of the Prime 2.0 analysis, which also takes into account the predicted strength of the HLA-peptide-TCR interaction. Estimated relative values ('ranking classes'; see 'Methods' for details) for the HLA/TCR interaction are shown on a background of decreasing grey color intensities, which, as explained in the scale caption, correspond to decreasing percentile group values (only the values for the best-scoring HLA alleles are reported). The sign and extent of variation of the mutated peptides scores with respect to the prototype peptides are reported in the last column, where green- and red-colored boxes indicate a score increase or decrease, corresponding to a gain or loss of T cell reactivity, respectively. The color shading code is the same as that utilized for IEDB analysis (see scale caption).

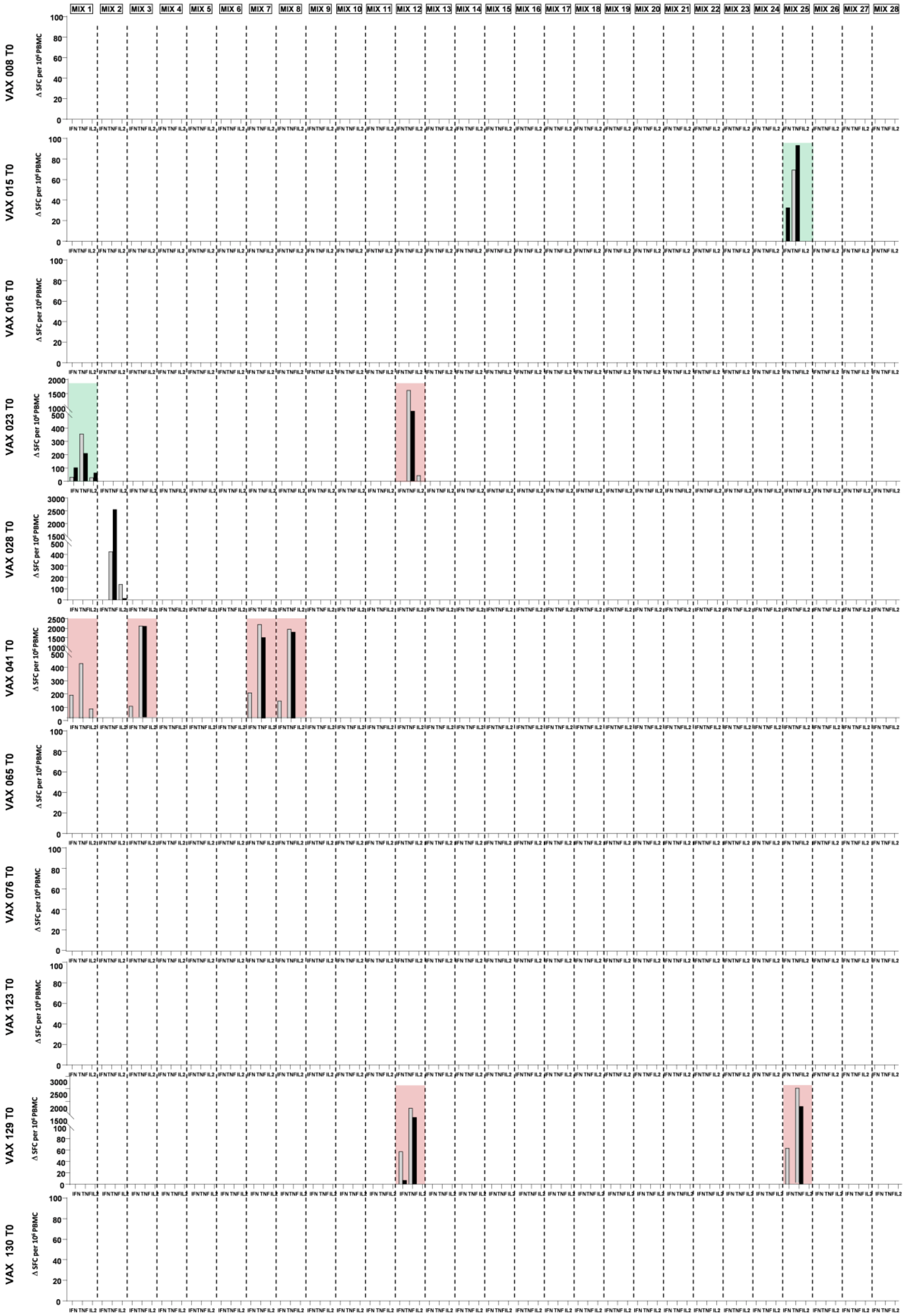

MIX 1
MIX 2
MIX 3
MIX 4
MIX 5
MIX 6
MIX 7
MIX 8
MIX 9
MIX 10
MIX 11
MIX 12
MIX 13
MIX 14
MIX 15
MIX 16
MIX 17
MIX 18
MIX 19
MIX 20
MIX 21
MIX 22
MIX 23
MIX 24
MIX 25
MIX 26
MIX 27
MIX 28
VAX 008 T0
VAX 015 T0
VAX 016 T0
VAX 023 T0
VAX 028 T0
VAX 041 T0
VAX 065 T0
VAX 076 T0
VAX 123 T0
VAX 129 T0
VAX 130 T0
Δ SFC per 10⁶ PBMC
IFN TNF IL2

**Extended Data Fig. 5. T cell responses in unvaccinated and uninfected individuals.** PBMCs from unvaccinated and uninfected individuals (n=11) were stimulated for 18 hours with overlapping wild-type and variant peptides. Cytokine-secreting T cells were measured by FluoroSPOT assays; all subjects are illustrated (see Fig. 1 legend for further details).

**Extended Data Table 1. Demographic and immunological parameters of the vaccine recipient population**

| | ID patient | Gender | Age | Days after 2nd vaccine dose | LOCUS A | LOCUS B | LOCUS C | SARS-CoV-2 serum neutralization assay after 2nd vaccine dose (NT50/mL) |
|---|---|---|---|---|---|---|---|---|
| **1** | **VAX 008*** | **F** | 43 | 17 | 11:01<br>24:02 | 51:01<br>- | 01:02<br>04:01 | ≥20480 |
| **2** | **VAX 009** | **F** | 62 | 46 | 01:01<br>03:01 | 35:02<br>40:02 | 02:02<br>04:01 | ≥20480 |
| **3** | **VAX 010** | **F** | 40 | 21 | 30:01<br>30:04 | 35:02<br>51:08 | 04:01<br>16:02 | ≥20480 |
| **4** | **VAX 011** | **M** | 48 | 39 | 02:01<br>26:01 | 18:01<br>35:01 | 04:01<br>07:01 | ≥5120 |
| **5** | **VAX 012** | **F** | 29 | 39 | 02:01<br>25:01 | 18:01<br>- | 07:01<br>12:03 | ≥20480 |
| **6** | **VAX 013** | **F** | 51 | 14 | 02:01<br>26:01 | 14:01<br>44:02 | 05:01<br>08:02 | ≥20480 |
| **7** | **VAX 015*** | **F** | 25 | 30 | 24:02<br>- | 18:01<br>44:02 | 02:02<br>12:03 | ≥20480 |
| **8** | **VAX 016*** | **F** | 28 | 30 | 02:01<br>32:01 | 18:01<br>40:02 | 02:02<br>07:01 | ≥20480 |
| **9** | **VAX 018** | **F** | 29 | 42 | 02:01<br>68:01 | 37:01<br>39:01 | 06:02<br>12:03 | ≥20480 |
| **10** | **VAX 019** | **F** | 51 | 23 | 03:01<br>33:01 | 13:02<br>14:02 | 07:04<br>08:02 | ≥20480 |
| **11** | **VAX 020** | **F** | 50 | 21 | 23:01<br>68:01 | 07:02<br>50:01 | 06:02<br>07:02 | ≥20480 |
| **12** | **VAX 021** | **F** | 53 | 38 | 02:01<br>11:01 | 15:17<br>51:01 | 07:01<br>07:01 | ≥20480 |
| **13** | **VAX 023*** | **F** | 49 | 29 | 01:01<br>24:02 | 45:01<br>52:01 | 07:02<br>12:02 | ≥10240 |
| **14** | **VAX 024** | **F** | 62 | 26 | 02:01<br>32:01 | 35:03<br>44:02 | 04:01<br>05:01 | ≥2560 |
| **15** | **VAX 025** | **M** | 37 | 41 | 02:01<br>26:26 | 13:02<br>27:05 | 02:02<br>06:02 | ≥20480 |
| **16** | **VAX 026** | **F** | 41 | 52 | 03:01<br>29:02 | 14:02<br>44:03 | 08:02<br>16:01 | ≥20480 |
| **17** | **VAX 027** | **M** | 48 | 29 | 02:01<br>30:02 | 18:01<br>39:01 | 05:01<br>12:03 | ≥20480 |
| **18** | **VAX 028*** | **F** | 39 | 69 | 01:01<br>26:01 | 08:01<br>38:01 | 07:01<br>12:03 | ≥10240 |
| **19** | **VAX 031** | **F** | 57 | 27 | 01:01<br>03:01 | 14:02<br>37:01 | 06:02<br>08:02 | ≥10240 |
| **20** | **VAX 033** | **F** | 54 | 27 | 02:01<br>30:02 | 35:02<br>44:02 | 04:01<br>05:01 | ≥20480 |
| **21** | **VAX 034** | **F** | 53 | 19 | 03:01<br>32:01 | 13:02<br>50:01 | 06:02<br>12:03 | ≥20480 |
| **22** | **VAX 035** | **F** | 57 | 26 | 30:01<br>33:01 | 13:02<br>14:02 | 06:02<br>08:02 | ≥20480 |
| **23** | **VAX 036** | **F** | 61 | 28 | 01:01<br>23:01 | 15:01<br>44:03 | 04:01<br>07:01 | ≥20480 |
| **24** | **VAX 039** | **F** | 62 | 23 | 02:01<br>03:01 | 44:02<br>57:01 | 05:01<br>07:01 | ≥20480 |
| **25** | **VAX 041*** | **F** | 27 | 15 | 03:01<br>24:02 | 07:02<br>51:01 | 07:02<br>15:02 | ≥20480 |
| **26** | **VAX 042** | **F** | 44 | 55 | 01:01<br>26:01 | 37:01<br>39:01 | 06:02<br>12:03 | ≥20480 |
| **27** | **VAX 043** | **F** | 50 | 31 | 01:01<br>02:01 | 27:02<br>38:01 | 02:02P<br>12:03P | ≥20480++++ |
| **28** | **VAX 044** | **M** | 65 | 38 | 01:01<br>26:01 | 08:01<br>44:05 | 02:02<br>07:01 | ≥20480 |
| **29** | **VAX 053** | **F** | 26 | 43 | 03:01<br>23:01 | 07:02<br>44:03 | 04:01<br>07:02 | ≥20480 |
| **30** | **VAX 055** | **F** | 27 | 58 | 11:01<br>30:01 | 13:02<br>35:01 | 04:01<br>06:02 | ≥20480 |
| **31** | **VAX 058** | **M** | 31 | 53 | 02:01<br>23:01P | 49:01<br>51:01 | 07:01<br>15:02 | ≥5120 |
| **32** | **VAX 060** | **F** | 28 | 59 | 02:01<br>03:01 | 35:02<br>51:01 | 04:01<br>16:02 | ≥20480 |
| **33** | **VAX 061** | **F** | 28 | 59 | 02:01<br>03:01 | 07:05<br>56:01 | 01:02<br>15:05 | ≥10240 |
| **34** | **VAX 065*** | **F** | 46 | 11 | 24:02<br>26:01 | 35:01<br>35:02 | 04:01<br>04:30 | ≥10240 |
| **35** | **VAX 066** | **M** | 58 | 58 | 02:05<br>11:01 | 35:01<br>58:01 | 04:01<br>07:18 | ≥2560 |
| **36** | **VAX 076*** | **F** | 26 | 31 | 03:01<br>24:02 | 07:02<br>35:08 | 04:01<br>07:02 | ≥10240 |

***** The asterix indicates the vaccinated subjects who have also been tested before vaccination, as a control for the stimulatory activity of the SARS-CoV-2 peptides in the absence of previous SARS-CoV-2 exposure. Three additional subjects, named as VAX 123, VAX 129 and VAX 130 have been enrolled to increase the number of control samples.

**Extended Data Table 2. VOC and VOI Spike mutations within each overlapping peptide pool**

| Domain | Wuhan-Hu1 | Alpha | Beta | Gamma | Delta | Epsilon | Eta | Omicron | MIX |
|---|---|---|---|---|---|---|---|---|---|
| | Mutation Site | B.1.1.7 | B.1.351 | P.1 | B.1.617.2 | B.1.429 | P.1.525 | B.1.1.529 | |
| 0 | S13 | | | | | I* | | | 25 |
| | L18 | | F | F | | | | | 1,2 |
| | T19 | | | | R** | | | | |
| | T20 | | | N | | | | | 1 |
| | P26 | | | S | | | | | 1 |
| | Q52 | | | | | | R | | 24 |
| | A67 | | | | | | | V | |
| | del 69-70 | del | | | | | del | del | 3 |
| | D80 | | A | | | | | | 4 |
| | T95 | | | | I | | | I | |
| | D138 | | | Y | | | | | 5 |
| S1 - NTD | W152 | | | | | C | | | 26 |
| | G142 | | | | D | | | D | |
| | del 143-145 | | | | | | | del | |
| | del 144 | del | | | | | | del | 6 |
| | E156 | | | | G | | | | |
| | Del 157-158 | | | | del | | | | |
| | R190 | | | S | | | | | 7 |
| | N211 | | | | | | | I | |
| | Del 212 | | | | | | | del | |
| | ins 214 | | | | | | | EPE | |
| | D215 | | G | | | | | | 8 |
| | del 242-244 | | del | | | | | | 9 |
| | R246 | | I | | | | | | 9 |
| 328 | G339 | | | | | | | D | |
| | S371 | | | | | | | L | |
| | S373 | | | | | | | P | |
| | S375 | | | | | | | F | |
| | K417 | | N | N | | | | N | 10 |
| | G446 | | | | | | | S | |
| S1 - RBD | L452 | | | | R | R | | | 27 |
| | S477 | | | | | | | N | |
| | T478 | | | | K | | | K | |
| | E484 | | K | K | | | K | A | 11 |
| | Q493 | | | | | | | R | |
| | G496 | | | | | | | S | |
| | Q498 | | | | | | | R | |
| | N501 | Y | Y | Y | | | | Y | 12 |
| | Y505 | | | | | | | H | |
| 533 | T547 | | | | | | | K | |
| | A570 | D | | | | | | | 13 |
| | D614 | G | G | G | G | G | G | G | 14 |
| S1-CTD1-2 | H655 | | | Y | | | | Y | 15 |
| | Q677 | | | | | | H | | 23 |
| | N679 | | | | | | | K | |
| | P681 | H | | | R | | | H | 16 |
| 686 | A701 | | V | | | | | | 17 |
| | T716 | I | | | | | | | 18 |
| | D796 | | | | | | | Y | |
| | N856 | | | | | | | K | |
| | F888 | | | | | | L | | 22 |
| | D950 | | | | N | | | | |
| S2 | Q954 | | | | | | | H | |
| | N969 | | | | | | | K | |
| | L981 | | | | | | | F | |
| | S982 | A | | | | | | | 19 |
| | T1027 | | | I | | | | | 20 |
| | D1118 | H | | | | | | | 21 |
| 1273 | V1176 | | | F | | | | | 28 |

* Red letters depict mutations contained in the overlapping peptide set used in the present study

** Black letters indicate mutations which have not been included in the peptide pools (most of them are expressed by the delta and omicron variants that have been reported when our study was already in progress)

**Extended Data Table 3. Overlapping wild type (Wuhan strain) and variant (VOC/VOI mutations) 10-mer peptides containing the VOC and VOI mutations illustrated in table S2.** Numbers in red indicate the position of the corresponding amino acid residue in the Spike sequence.

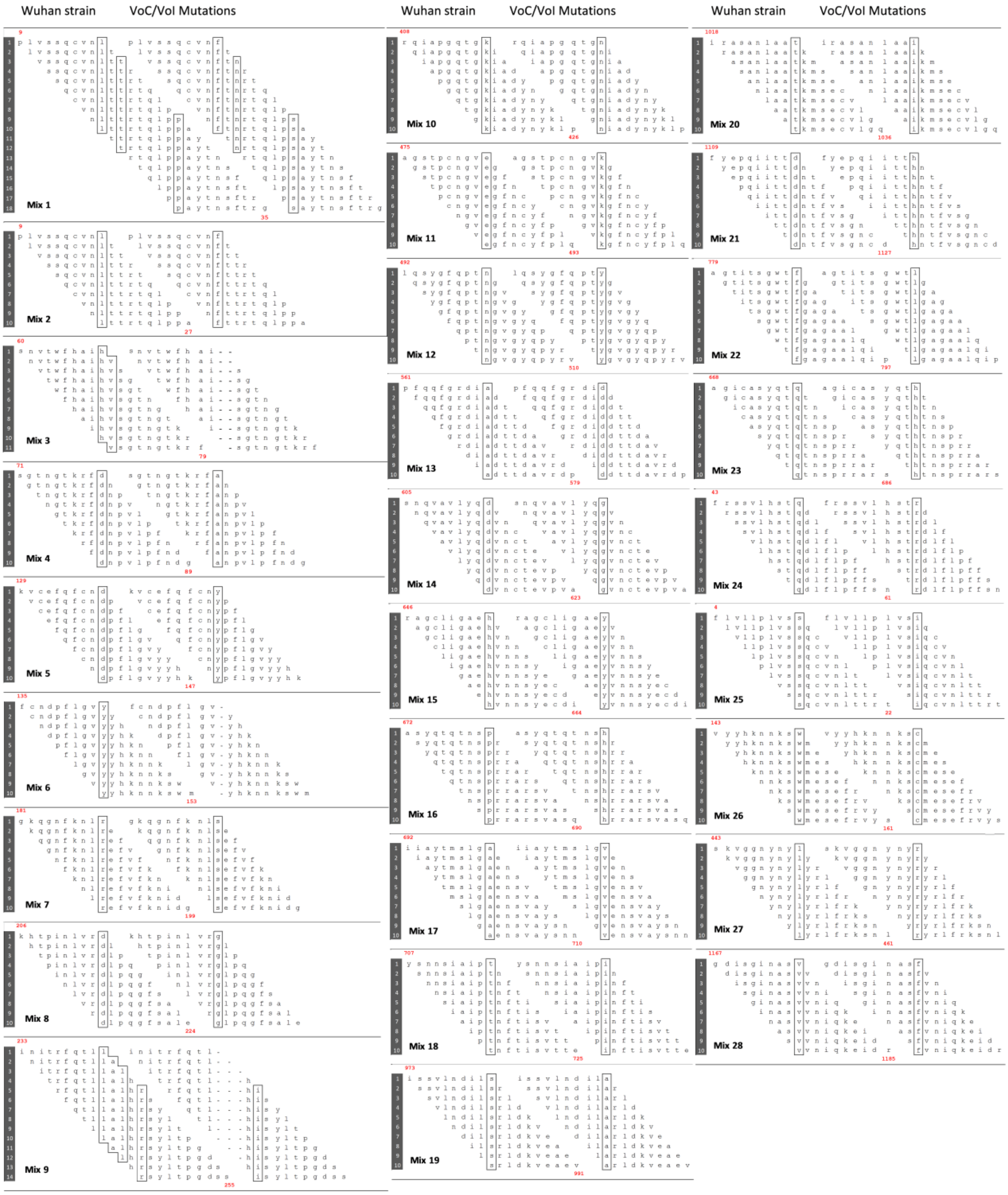

**Extended Data Table 4. Demographic and clinical features of the recruited convalescent patients**

| | ID patient | Gender | Age | Days elapsed between the first positive swab and the immunological sample | Lung tomography (visual score %) | Steroid therapy (Y/N) | Oxygen therapy (Y/N) | Non-invasive ventilation (Y/N) | C-reactive protein | D-dimer | White blood cells | Relevant comorbidities |
|---|---|---|---|---|---|---|---|---|---|---|---|---|
| 1 | COV 001 | F | 60 | 35 | 35% | N (tocilizumab) | Y | Y | 136 mg/L | 898 ng/mL | 3950 x $10^3$/mL | |
| 2 | COV 002 | M | 64 | 48 | severe: 70% | Y | Y | Y | > 250 mg/L | 5573 ng/mL | 20620 x $10^3$/mL | |
| 3 | COV 003 | F | 66 | 11 | 40% | N | Y | N | > 79 mg/L | 1348 ng/mL | 4160 x $10^3$/mL | |
| 4 | COV 008 | M | 55 | 21 | severe: 70% | Y | Y | Y | 137 mg/L | 885 ng/mL | 7560 x $10^3$/mL | |
| 5 | COV 009 | F | 61 | 42 | severe: 45% | Y + tocilizumab | Y | Y + IOT* | > 250 mg/L | 560 ng/mL --> 9000** | 7920 x $10^3$/mL | obesity |
| 6 | COV 010 | M | 45 | 18 | 20% | Y | Y | Y | 240 mg/L | 2510 ng/mL | 18530 x $10^3$/mL | kidney tumor |
| 7 | COV 011 | M | 57 | 46 | severe: 60% | N | Y | Y | 246 mg/L | 2301 ng/mL | 6140 x $10^3$/mL | pneumothorax |
| 8 | COV 012 | F | 77 | 44 | severe: 60% | Y + tocilizumab | Y | Y | 65 mg/L | > 9000 ng/mL | 8060 x $10^3$/mL | |
| 9 | COV 014 | F | 68 | 6 | 0% | N | N | N | 52 mg/L | 1085 ng/mL | 3550 x $10^3$/mL | lung cancer and recent chemotherapy |
| 10 | COV 017 | M | 58 | 12 | 25% | N | N | N | 4.8 mg/L | 641 ng/mL | 4890 x $10^3$/mL | |
| 11 | COV 022 | M | 46 | 17 | 10% | N | N | N | 19 mg/L | 400 ng/mL | 5420 x $10^3$/mL | |
| 12 | COV 024 | F | 65 | 14 | neg | N | N | N | 99 mg/L | 3853 ng/mL | 11580 x $10^3$/mL | |

IOT*: tracheal intubation
** peripheral venous thrombosis

**Extended Data Table 5. Homology between enhancing SARS-CoV-2 mutated peptides and pre-existing human pathogen sequences.** Red letters depict the different amino acids present in the homologous peptides with respect to those present in the enhancing mutated peptides. The boxed letters or vertical lines highlight the location of VOC and VOI amino acid changes or deletions, respectively.

| FCNYPFLGVY | Organism | Accession number | Sequence | Identity | Similarity | Taxonomy |
|---|---|---|---|---|---|---|
| | SarsCov2(mut) | | F C N Y P F L G V Y | 10 | 10 | V |
| | Burkholderia cenocepacia | WP_084750340.1 | F C G Y P G L G V Y | 8 | 8 | B |
| | Chloroflexi bacterium | MBI5960450.1 | F A N Y P F L G V T | 8 | 8 | B |
| | Lachnospiraceae bacterium | MCD8397397.1 | F C N Y I F L G V K | 8 | 8 | B |
| | Bacteroidales bacterium | MBD5171519.1 | V Y N Y P F L G V Y | 8 | 8 | B |
| | Trichuris trichiura | CDW52749.1 | F C K N P F L G V Y | 8 | 8 | A |

P=plant
B=bacteria
V=virus
F=fungus
A=animal
U=protozoa

| GVYHKNNKSW | Organism | Accession number | Sequence | Identity | Similarity | Taxonomy |
|---|---|---|---|---|---|---|
| | SarsCov2(mut) | | G V Y H K N N K S W | 10 | 10 | V |
| | Avrilella dinanensis | WP_100677051.1 | G V Y H Q N N K S M | 8 | 8 | B |
| | Ruminococcus sp. | MBQ4247073.1 | G V Y H T N N K S T | 8 | 8 | B |
| | Bacteroidetes bacterium | MBT4337516.1 | G V Y K K N N K S I | 8 | 9 | B |
| | Trypanosoma theileri | XP_028885404.1 | G V Y H K I N K R W | 8 | 8 | U |
| | Lachnospiraceae bacterium | MBR1440785.1 | G V Y H K N N T S R | 8 | 8 | B |
| | Roseburia sp. | WP_022516031.1 | G V Y H K N N S S R | 8 | 8 | B |
| | Clostridium sp. | PWM81222.1 | G V Y F K N N K S T | 8 | 8 | B |
| | Campylobacter lari | EAK6287402.1 | G V Y S K N N K S Y | 8 | 8 | B |
| | Elusimicrobia bacterium | OGS37176.1 | G V Y N K N N K S A | 8 | 8 | B |

| VKGFNCYFPL | Organism | Accession number | Sequence | Identity | Similarity | Taxonomy |
|---|---|---|---|---|---|---|
| | SarsCov2(mut) | | V K G F N C Y F P L | 10 | 10 | V |
| | Bacteroidia bacterium | MBI4648754.1 | V K G F S C Y F G L | 8 | 8 | B |
| | Verrucomicrobia bacterium | RKX37078.1 | G K G F N C Y F P M | 8 | 8 | B |
| | Firmicutes bacterium | HFT74624.1 | D K G F N C Y F P M | 8 | 8 | B |
| | Proteobacteria bacterium | MBE9545667.1 | G K G F N C Y F P M | 8 | 8 | B |

| IPINFTISVT | Organism | Accession number | Sequence | Identity | Similarity | Taxonomy |
|---|---|---|---|---|---|---|
| | SarsCov2(mut) | | I P I N F T I S V T | 10 | 10 | V |
| | Rickettsiales bacterium | MCE2926686.1 | I P I N F G I S V T | 9 | 9 | B |
| | Clostridiales bacterium | KXO15851.1 | I P I - F T I S V T | 9 | 9 | B |
| | Gram-Positive Anaerobic Cocci | WP_068367975.1<br>WP_044566338.1<br>MBS6611571.1<br>WP_004167031.1<br>WP_202947903.1 | I P I - F T I S V T | 9 | 9 | B |
| | Enterococcus faecium | HBM7176442.1 | I P I - F T I S V T | 9 | 9 | B |
| | Paeniclostridium sordellii | MBS6025935.1 | I P I - F T I S V T | 9 | 9 | B |
| | Prevotella koreensis | WP_126678470.1 | V P I N F T I N V T | 8 | 9 | B |
| | Clostridiales bacterium | NLO47780.1 | I S V N F T I S V T | 8 | 9 | B |
| | | MBQ8556355.1 | I N V N F T I S V T | 8 | 9 | B |
| | Limosilactobacillus antri | WP_007124064.1 | I P V N F T I S V A | 8 | 9 | B |
| | Archaeon | NAT11278.1 | M P I N F T V S V T | 8 | 9 | B |
| | Nocardia sp. | WP_228001339.1 | I G I D F T I S V T | 8 | 8 | B |
| | Bacteroidaceae bacterium | MBR6856320.1 | I P I N I T F S V T | 8 | 8 | B |
| | Burkholderiales bacterium | MBC7784469.1 | I P I N F G I S V V | 8 | 8 | B |
| | Proteobacteria bacterium | MBS0185753.1 | I P I N F G I S V F | 8 | 8 | B |
| | Rickettsiaceae bacterium | NRB10244.1 | I P I N F G I S V L | 8 | 8 | B |
| | Gammaproteobacteria bacterium | NBS15877.1 | I P I N F G I S V A | 8 | 8 | B |
| | Chlamydiae bacterium | MBM3197346.1 | I P I N F G I S V L | 8 | 8 | B |
| | Trichomonas vaginalis G3 | XP_001321750.1 | I P I N F T I S S E | 8 | 8 | U |
| | Loigolactobacillus bifermentans | WP_057905577.1 | Q P I N F T I S V G | 8 | 8 | B |
| | Firmicutes | WP_117994809.1 | I P E N F T I S V F | 8 | 8 | B |
| | Chlamydiales bacterium | NCF70221.1 | I P E N F T I S V D | 8 | 8 | B |

| PLVSIQCVNL | Organism | Accession number | Sequence | Identity | Similarity | Taxonomy |
|---|---|---|---|---|---|---|
| | SarsCov2(mut) | | P L V S I Q C V N L | 10 | 10 | V |
| | Rhizopus arrhizus | KAG1069288.1 | P F V S I Q C Q N L | 8 | 8 | F |
| | Dysgonomonas massiliensis | WP_245871973.1 | F L V S I L C V N L | 8 | 8 | B |
| | Dysgonomonas sp. 521 | WP_163308040.1 | F L V S I I C V N L | 8 | 8 | B |
| | Rikenellaceae bacterium | MBP3425517.1 | E L V S I Q C V N D | 8 | 8 | B |

| LVLLPLVSIQ | Organism | Accession number | Sequence | Identity | Similarity | Taxonomy |
|---|---|---|---|---|---|---|
| | SarsCov2(mut) | | L V L L P L V S I Q | 10 | 10 | V |
| | Aspergillus versicolor | XP_040663065.1 | L L L L P L V S I Q | 9 | 10 | F |
| | Aspergillus sp. | RMJ24681.1 | L L L L P L V S I Q | 9 | 10 | F |
| | Burkholderiaceae bacterium | MBW6496305.1 | L V L L P L V R I Q | 9 | 9 | B |
| | Lachnospiraceae bacterium | MBD5508727.1 | L V L L L L V S I Q | 9 | 9 | B |
| | Vibrio sp. | WP_223100057.1 | L V L L P L V S I A | 9 | 9 | B |
| | Pseudomonadales bacterium | MBL6687889.1 | L I L L A L V S I Q | 8 | 9 | B |
| | Clostridia bacterium | MBQ8093389.1<br>MBS7141494.1 | L V L L P L I S I I/L | 8 | 9 | B |
| | Clostridium sp. | WP_176255526.1 | L V L L P L I S I L | 8 | 9 | B |
| | Spirochaetae bacterium | PKL18185.1 | L V L L P L V S V W | 8 | 9 | B |
| | Actinomyces sp. | WP_223911791.1<br>WP_231945305.1 | L V L L P L V T I L | 8 | 9 | B |
| | Clostridia bacterium | MBR4186193.1 | L V L L P L V S V I | 8 | 9 | B |
| | Mycobacterium sp. | MBB5165227.1 | L V L L P L V S V S | 8 | 9 | B |
| | Bacteroidales bacterium | NLA64047.1 | L V L L P L V T I I | 8 | 9 | B |
| | Oscillospiraceae bacterium | MBE6541461.1 | L V L L P I V S I I | 8 | 9 | B |
| | Gammaproteobacteria bacterium | OGT65810.1 | L V L L P L V S V S | 8 | 9 | B |

| SIQCVNLTTR | Organism | Accession number | Sequence | Identity | Similarity | Taxonomy |
|---|---|---|---|---|---|---|
| | SarsCov2(mut) | | S I Q C V N L T T R | 10 | 10 | V |
| | Bacteroides sp. | WP_181981949.1<br>MBS5410381.1<br>AAO78114.1 | S I H C V N L P T R | 8 | 8 | B |
| | Clostridia bacterium | NCB74989.1 | S I Q C V N L N T L | 8 | 8 | B |
| | Aspergillus nidulans | XP_680781.1 | D I Q C V I L T T R | 8 | 8 | F |
| | Loa loa | XP_003137001.1 | S I Q S V N Q T T R | 8 | 8 | A |

| PELGVYHKNN | Organism | Accession number | Sequence | Identity | Similarity | Taxonomy |
|---|---|---|---|---|---|---|
| | SarsCov2(mut) | | P F L G V Y H K N N | 10 | 10 | V |
| | Bacteroides sp. | MCD8260604.1 | P Y L G V Y H K N N | 9 | 10 | B |
| | Bacteroidetes bacterium | MBK7440535.1 | P F L E V Y R K N N | 8 | 9 | B |
| | Mycobacteriaceae bacterium | MBV9089423.1<br>WP_030136828.1<br>BBY44688.1<br>WP_064872347.1<br>WP_085192734.1<br>MBU8829181.1<br>WP_058125021.1<br>WP_067392814.1<br>WP_064399970.1<br>KUI05088.1<br>WP_094292683.1 | P F L G V Y R K A N | 8 | 9 | B |
| | Bacilli bacterium | MBQ3475877.1 | P F L G V Y R K A N | 8 | 9 | B |
| | Trichomonas vaginalis | XP_001321691.1 | P F L M A Y H K N N | 8 | 8 | U |
| | Clostridia bacterium | MBQ9781832.1 | P F L G E Y H K N G | 8 | 8 | B |
| | Ruminococcus sp. | MCC8192177.1<br>MCD7726859.1 | P F L - V Y H K N G | 8 | 8 | B |
| | Mycolicibacterium sp. | WP_207359507.1<br>WP_005142699.1<br>WP_069412734.1 | P F L G V Y R D N N | 8 | 9 | B |
| | Lachnospiraceae bacterium | MCI8453462.1 | R F L G V Y E K N N | 8 | 8 | B |
| | Clostridia bacterium | MBQ8294862.1 | P F L G V Y N N N T | 8 | 8 | B |
| | Clostridia bacterium | MBQ8320281.1 | P F L G V K H K D N | 8 | 8 | B |

| LPLVSIQCVN | Organism | Accession number | Sequence | Identity | Similarity | Taxonomy |
|---|---|---|---|---|---|---|
| | SarsCov2(mut) | | L P L V S I Q C V N | 10 | 10 | V |
| | Clostridia bacterium | MBR0414238.1 | L Q V V S I Q C V N | 8 | 9 | B |
| | Oscillospiraceae bacterium | MBE6819079.1 | L Q V V S I Q C V N | 8 | 9 | B |
| | Bacteroidaceae bacterium | MBO7291606.1 | D P L V S I Q C I N | 8 | 9 | B |
| | Bacteroidales bacterium | MBN1989265.1 | L P L V A I Q V V N | 8 | 8 | B |
| | Arachnia propionica | WP_061787963.1 | L P L V S I S C V S | 8 | 8 | B |
| | Slackia piriformis | WP_009139958.1 | L P L V S I L C V Y | 8 | 8 | B |
| | Propionibacterium sp. | MBB1587979.1 | L P L V S I S C V S | 8 | 8 | B |
| | Firmicutes bacterium | PKM39155.1 | L P L V S I G C V M | 8 | 8 | B |
| | Clostridiales bacterium | NLI54870.1 | L P L V S I G C V V | 8 | 8 | B |
| | Actinomycetia bacterium | MTA82618.1 | L P L V S I V C V V | 8 | 8 | B |
| | Rikenellaceae bacterium | MBP3425517.1 | H E L V S I Q C V N | 8 | 8 | B |
| | Prevotella koreensis | WP_126678039.1 | L P L H S M Q C V N | 8 | 8 | B |
| | Trichinella nelsoni | KRX26147.1 | L P L V S P Q C V L | 8 | 8 | B |

**Extended Data Table 6. List of overlapping 15 mer peptides spanning the entire spike sequence**

| | POSITION | SEQUENCE |
|---|---|---|
| S1 pool (34 peptides) | 1-15 | MFVFLVLLPLVSSQC |
| | 6-20 | VLLPLVSSQCVNLTT |
| | 11-25 | VSSQCVNLTTRTQLP |
| | 16-30 | VNLTTRTQLPPAYTN |
| | 21-35 | RTQLPPAYTNSFTRG |
| | 26-40 | PAYTNSFTRGVYYPD |
| | 31-45 | SFTRGVYYPDKVFRS |
| | 36-50 | VYYPDKVFRSSVLHS |
| | 41-55 | KVFRSSVLHSTQDLF |
| | 46-60 | SVLHSTQDLFLPFFS |
| | 51-65 | TQDLFLPFFSNVTWF |
| | 56-70 | LPFFSNVTWFHAIHV |
| | 61-75 | NVTWFHAIHVSGTNG |
| | 66-80 | HAIHVSGTNGTKRFD |
| | 71-85 | SGTNGTKRFDNPVLP |
| | 76-90 | TKRFDNPVLPFNDGV |
| | 81-95 | NPVLPFNDGVYFAST |
| | 86-100 | FNDGVYFASTEKSNI |
| | 91-105 | YFASTEKSNIIRGWI |
| | 96-110 | EKSNIIRGWIFGTTL |
| | 101-115 | IRGWIFGTTLDSKTQ |
| | 106-120 | FGTTLDSKTQSLLIV |
| | 111-125 | DSKTQSLLIVNNATN |
| | 116-130 | SLLIVNNATNVVIKV |
| | 121-135 | NNATNVVIKVCEFQF |
| | 126-140 | VVIKVCEFQFCNDPF |
| | 131-145 | CEFQFCNDPFLGVYY |
| | 136-150 | CNDPFLGVYYHKNNK |
| | 141-155 | LGVYYHKNNKSWMES |
| | 146-160 | HKNNKSWMESEFRVY |
| | 151-165 | SWMESEFRVYSSANN |
| | 156-170 | EFRVYSSANNCTFEY |
| | 161-175 | SSANNCTFEYVSQPF |
| | 166-180 | CTFEYVSQPFLMDLE |
| S2 pool (33 peptides) | 171-185 | VSQPFLMDLEGKQGN |
| | 176-190 | LMDLEGKQGNFKNLR |
| | 181-195 | GKQGNFKNLREFVFK |
| | 186-200 | FKNLREFVFKNIDGY |
| | 191-205 | EFVFKNIDGYFKIYS |
| | 196-210 | NIDGYFKIYSKHTPI |
| | 201-215 | FKIYSKHTPINLVRD |
| | 206-220 | KHTPINLVRDLPQGF |
| | 211-225 | NLVRDLPQGFSALEP |
| | 216-230 | LPQGFSALEPLVDLP |
| | 221-235 | SALEPLVDLPIGINI |
| | 226-240 | LVDLPIGINITRFQT |
| | 231-245 | IGINITRFQTLLALH |
| | 236-250 | TRFQTLLALHRSYLT |
| | 241-255 | LLALHRSYLTPGDSS |
| | 246-260 | RSYLTPGDSSSGWTA |
| | 251-265 | PGDSSSGWTAGAAAY |
| | 256-270 | SGWTAGAAAYYVGYL |
| | 261-275 | GAAAYYVGYLQPRTF |
| | 266-280 | YVGYLQPRTFLLKYN |
| | 271-285 | QPRTFLLKYNENGTI |
| | 276-290 | LLKYNENGTITDAVD |
| | 281-295 | ENGTITDAVDCALDP |
| | 286-300 | TDAVDCALDPLSETK |
| | 291-305 | CALDPLSETKCTLKS |
| | 296-310 | LSETKCTLKSFTVEK |
| | 301-315 | CTLKSFTVEKGIYQT |
| | 306-320 | FTVEKGIYQTSNFRV |
| | 311-325 | GIYQTSNFRVQPTES |
| | 316-330 | SNFRVQPTESIVRFP |
| | 321-335 | QPTESIVRFPNITNL |
| | 326-340 | IVRFPNITNLCPFGE |
| | 331-345 | NITNLCPFGEVFNAT |
| S3 pool (33 peptides) | 336-350 | CPFGEVFNATRFASV |
| | 341-355 | VFNATRFASVYAWNR |
| | 346-360 | RFASVYAWNRKRISN |
| | 351-365 | YAWNRKRISNCVADY |
| | 356-370 | KRISNCVADYSVLYN |
| | 361-375 | CVADYSVLYNSASFS |
| | 366-380 | SVLYNSASFSTFKCY |
| | 371-385 | SASFSTFKCYGVSPT |
| | 376-390 | TFKCYGVSPTKLNDL |
| | 381-395 | GVSPTKLNDLCFTNV |
| | 386-400 | KLNDLCFTNVYADSF |
| | 391-405 | CFTNVYADSFVIRGD |
| | 396-410 | YADSFVIRGDEVRQI |
| | 401-415 | VIRGDEVRQIAPGQT |
| | 406-420 | EVRQIAPGQTGKIAD |
| | 411-425 | APGQTGKIADYNYKL |
| | 416-430 | GKIADYNYKLPDDFT |
| | 421-435 | YNYKLPDDFTGCVIA |
| | 426-440 | PDDFTGCVIAWNSNN |
| | 431-445 | GCVIAWNSNNLDSKV |
| | 436-450 | WNSNNLDSKVGGNYN |
| | 441-455 | LDSKVGGNYNYLYRL |
| | 446-460 | GGNYNYLYRLFRKSN |
| | 451-465 | YLYRLFRKSNLKPFE |
| | 456-470 | FRKSNLKPFERDIST |
| | 461-475 | LKPFERDISTEIYQA |
| | 466-480 | RDISTEIYQAGSTPC |
| | 471-485 | EIYQAGSTPCNGVEG |
| | 476-490 | GSTPCNGVEGFNCYF |
| | 481-495 | NGVEGFNCYFPLQSY |
| | 486-500 | FNCYFPLQSYGFQPT |
| | 491-505 | PLQSYGFQPTNGVGY |
| | 496-510 | GFQPTNGVGYQPYRV |
| S4 pool (39 peptides) | 501-515 | NGVGYQPYRVVVLSF |
| | 506-520 | QPYRVVVLSFELLHA |
| | 511-525 | VVLSFELLHAPATVC |
| | 516-530 | ELLHAPATVCGPKKS |
| | 521-535 | PATVCGPKKSTNLVK |
| | 526-540 | GPKKSTNLVKNKCVN |
| | 531-545 | TNLVKNKCVNFNFNG |
| | 536-550 | NKCVNFNFNGLTGTG |
| | 541-555 | FNFNGLTGTGVLTES |
| | 546-560 | LTGTGVLTESNKKFL |
| | 551-565 | VLTESNKKFLPFQQF |
| | 556-570 | NKKFLPFQQFGRDIA |
| | 561-575 | PFQQFGRDIADTTDA |
| | 566-580 | GRDIADTTDAVRDPQ |
| | 571-585 | DTTDAVRDPQTLEIL |
| | 576-590 | VRDPQTLEILDITPC |
| | 581-595 | TLEILDITPCSFGGV |
| | 586-600 | DITPCSFGGVSVITP |
| | 591-605 | SFGGVSVITPGTNTS |
| | 596-610 | SVITPGTNTSNQVAV |
| | 601-615 | GTNTSNQVAVLYQDV |
| | 606-620 | NQVAVLYQDVNCTEV |
| | 611-625 | LYQDVNCTEVPVAIH |
| | 616-630 | NCTEVPVAIHADQLT |
| | 621-635 | PVAIHADQLTPTWRV |
| | 626-640 | ADQLTPTWRVYSTGS |
| | 631-645 | PTWRVYSTGSNVFQT |
| | 636-650 | YSTGSNVFQTRAGCL |
| | 641-655 | NVFQTRAGCLIGAEH |
| | 646-660 | RAGCLIGAEHVNNSY |
| | 651-665 | IGAEHVNNSYECDIP |
| | 656-670 | VNNSYECDIPIGAGI |
| | 661-675 | ECDIPIGAGICASYQ |
| | 666-680 | IGAGICASYQTQTNS |
| | 671-685 | CASYQTQTNSPRRAR |
| | 676-690 | TQTNSPRRARSVASQ |
| | 681-695 | PRRARSVASQSIIAY |
| | 686-700 | SVASQSIIAYTMSLG |
| | 691-705 | SIIAYTMSLGAENSV |
| S5 pool (38 peptides) | 696-710 | TMSLGAENSVAYSNN |
| | 701-715 | AENSVAYSNNSIAIP |
| | 706-720 | AYSNNSIAIPTNFTI |
| | 711-725 | SIAIPTNFTISVTTE |
| | 716-730 | TNFTISVTTEILPVS |
| | 721-735 | SVTTEILPVSMTKTS |
| | 726-740 | ILPVSMTKTSVDCTM |
| | 731-745 | MTKTSVDCTMYICGD |
| | 736-750 | VDCTMYICGDSTECS |
| | 741-755 | YICGDSTECSNLLLQ |
| | 746-760 | STECSNLLLQYGSFC |
| | 751-765 | NLLLQYGSFCTQLNR |
| | 756-770 | YGSFCTQLNRALTGI |
| | 761-775 | TQLNRALTGIAVEQD |
| | 766-780 | ALTGIAVEQDKNTQE |
| | 771-785 | AVEQDKNTQEVFAQV |
| | 776-790 | KNTQEVFAQVKQIYK |
| | 781-795 | VFAQVKQIYKTPPIK |
| | 786-800 | KQIYKTPPIKDFGGF |
| | 791-805 | TPPIKDFGGFNFSQI |
| | 796-810 | DFGGFNFSQILPDPS |
| | 801-815 | NFSQILPDPSKPSKR |
| | 806-820 | LPDPSKPSKRSFIED |
| | 811-825 | KPSKRSFIEDLLFNK |
| | 816-830 | SFIEDLLFNKVTLAD |
| | 821-835 | LLFNKVTLADAGFIK |
| | 826-840 | VTLADAGFIKQYGDC |
| | 831-845 | AGFIKQYGDCLGDIA |
| | 836-850 | QYGDCLGDIAARDLI |
| | 841-855 | LGDIAARDLICAQKF |
| | 846-860 | ARDLICAQKFNGLTV |
| | 851-865 | CAQKFNGLTVLPPLL |
| | 856-870 | NGLTVLPPLLTDEMI |
| | 861-875 | LPPLLTDEMIAQYTS |
| | 866-880 | TDEMIAQYTSALLAG |
| | 871-885 | AQYTSALLAGTITSG |
| | 876-890 | ALLAGTITSGWTFGA |
| | 881-895 | TITSGWTFGAGAALQ |
| S6 pool (38 peptides) | 886-900 | WTFGAGAALQIPFAM |
| | 891-905 | GAALQIPFAMQMAYR |
| | 896-910 | IPFAMQMAYRFNGIG |
| | 901-915 | QMAYRFNGIGVTQNV |
| | 906-920 | FNGIGVTQNVLYENQ |
| | 911-925 | VTQNVLYENQKLIAN |
| | 916-930 | LYENQKLIANQFNSA |
| | 921-935 | KLIANQFNSAIGKIQ |
| | 926-940 | QFNSAIGKIQDSLSS |
| | 931-945 | IGKIQDSLSSTASAL |
| | 936-950 | DSLSSTASALGKLQD |
| | 941-955 | TASALGKLQDVVNQN |
| | 946-960 | GKLQDVVNQNAQALN |
| | 951-965 | VVNQNAQALNTLVKQ |
| | 956-970 | AQALNTLVKQLSSNF |
| | 961-975 | TLVKQLSSNFGAISS |
| | 966-980 | LSSNFGAISSVLNDI |
| | 971-985 | GAISSVLNDILSRLD |
| | 976-990 | VLNDILSRLDKVEAE |
| | 981-995 | LSRLDKVEAEVQIDR |
| | 986-1000 | KVEAEVQIDRLITGR |
| | 991-1005 | VQIDRLITGRLQSLQ |
| | 996-1010 | LITGRLQSLQTYVTQ |
| | 1001-1015 | LQSLQTYVTQQLIRA |
| | 1006-1020 | TYVTQQLIRAAEIRA |
| | 1011-1025 | QLIRAAEIRASANLA |
| | 1016-1030 | AEIRASANLAATKMS |
| | 1021-1035 | SANLAATKMSECVLG |
| | 1026-1040 | ATKMSECVLGQSKRV |
| | 1031-1045 | ECVLGQSKRVDFCGK |
| | 1036-1050 | QSKRVDFCGKGYHLM |
| | 1041-1055 | DFCGKGYHLMSFPQS |
| | 1046-1060 | GYHLMSFPQSAPHGV |
| | 1051-1065 | SFPQSAPHGVVFLHV |
| | 1056-1070 | APHGVVFLHVTYVPA |
| | 1061-1075 | VFLHVTYVPAQEKNF |
| | 1066-1080 | TYVPAQEKNFTTAPA |
| | 1071-1085 | QEKNFTTAPAICHDG |
| S7 pool (38 peptides) | 1076-1090 | TTAPAICHDGKAHFP |
| | 1081-1095 | ICHDGKAHFPREGVF |
| | 1086-1100 | KAHFPREGVFVSNGT |
| | 1091-1105 | REGVFVSNGTHWFVT |
| | 1096-1110 | VSNGTHWFVTQRNFY |
| | 1101-1115 | HWFVTQRNFYEPQII |
| | 1106-1120 | QRNFYEPQIITTDNT |
| | 1111-1125 | EPQIITTDNTFVSGN |
| | 1116-1130 | TTDNTFVSGNCDVVI |
| | 1121-1135 | FVSGNCDVVIGIVNN |
| | 1126-1140 | CDVVIGIVNNTVYDP |
| | 1131-1145 | GIVNNTVYDPLQPEL |
| | 1136-1150 | TVYDPLQPELDSFKE |
| | 1141-1155 | LQPELDSFKEELDKY |
| | 1146-1160 | DSFKEELDKYFKNHT |
| | 1151-1165 | ELDKYFKNHTSPDVD |
| | 1156-1170 | FKNHTSPDVDLGDIS |
| | 1161-1175 | SPDVDLGDISGINAS |
| | 1166-1180 | LGDISGINASVVNIQ |
| | 1171-1185 | GINASVVNIQKEIDR |
| | 1176-1190 | VVNIQKEIDRLNEVA |
| | 1181-1195 | KEIDRLNEVAKNLNE |
| | 1186-1200 | LNEVAKNLNESLIDL |
| | 1191-1205 | KNLNESLIDLQELGK |
| | 1196-1210 | SLIDLQELGKYEQYI |
| | 1201-1215 | QELGKYEQYIKWPWY |
| | 1206-1220 | YEQYIKWPWYIWLGF |
| | 1211-1225 | KWPWYIWLGFIAGLI |
| | 1216-1230 | IWLGFIAGLIAIVMV |
| | 1221-1235 | IAGLIAIVMVTIMLC |
| | 1226-1240 | AIVMVTIMLCCMTSC |
| | 1231-1245 | TIMLCCMTSCCSCLK |
| | 1236-1250 | CMTSCCSCLKGCCSC |
| | 1241-1255 | CSCLKGCCSCGSCCK |
| | 1246-1260 | GCCSCGSCCKFDEDD |
| | 1251-1265 | GSCCKFDEDDSEPVL |
| | 1256-1270 | FDEDDSEPVLKGVKL |
| | 1261-1273 | SEPVLKGVKLHYT |

**Extended Data Table 7. List of the peptides reported in the literature and used to map CD8 T cell responses with indication of conservation and degeneracy levels**

| | Position | Sequence | C | HLA-Restricted | D | Reference |
|---|---|---|---|---|---|---|
| **L1 pool (11 peptides)** | 28-37 | YTNSFTRGVY | | **A*01** - **A*26** - A*30 - B*15 | * | 53 |
| | 35-44 | GVYYPDKVFR | | **A*68** - **A*03** - A*31 - A*11 | * | 53 |
| | 41-49 | KVFRSSVLH | | **A*30** - **A*03** - A*11 | * | 53 |
| | 109-117 | TLDSKTQSL | | **A*02** - **B*08** | * | 53 |
| | 118-126 | LIVNNATNV | | A*02 | | IEDB prediction tool |
| | 118-126 | LIVNDATNV | | A*02 | | IEDB prediction tool |
| | 269-277 | YLQPRTFLL | | **A*02** - **B*08** - A*23 - A*32 | * | 30, 31, 52, 53 |
| | 276-285 | LLKYNENGTI | | A*02 | | IEDB prediction tool |
| | 276-285 | LLKYNEDGTI | | A*02 | | IEDB prediction tool |
| | 285-293 | ITDAVDCAL | | A*02 | | IEDB prediction tool |
| | 292-300 | ALDPLSETK | | A*03 | | 53 |
| **L2 pool (11 peptides)** | 297-306 | SETKCTLKSF | | **B*44** | * | 53 |
| | 312-320 | IYQTSNFRV | | **A*24** - A*23 | | 53 |
| | 321-329 | QPTESIVRF | | **B*35** - **B*07** - B*53 | * | 53 |
| | 323-335 | TESIVRFPNITNL | | B*40 | | 53 |
| | 323-335 | TESIVRFPDITNL | | B*40 | | 53 |
| | 328-338 | RFPNITNLCPF | | **A*24** - B*35 - A*23 | | 53 |
| | 328-338 | RFPDITNLCPF | | **A*24** | | 53 |
| | 340-351 | EVFNATRFASVY | | **A*26** | | 53 |
| | 340-351 | EVFDATRFASVY | | **A*26** | | 53 |
| | 386-395 | KLNDLCFTNV | | **A*02** | | IEDB prediction tool |
| | 394-403 | NVYADSFVIR | | **A*68** - A*33 - A*31 | | 53 |
| **L3 pool (11 peptides)** | 400-408 | FVIRGDEVR | | **A*68** | | 53 |
| | 424-433 | KLPDDFTGCV | | **A*02** | | 30 |
| | 433-441 | VIAWNSNNL | | A*02 | | IEDB prediction tool |
| | 458-466 | KSNLKPFER | | **A*31** | | 53 |
| | 462-472 | KPFERDISTEI | | **B*07** | | 53 |
| | 506-515 | QPYRVVVLSF | | **B*07** - A*23 - A*24 | * | 53 |
| | 512-520 | VLSFELLHA | | A*02 | | IEDB prediction tool |
| | 526-534 | GPKKSTNLV | | B*07 | | IEDB prediction tool |
| | 529-537 | KSTNLVKNK | | A*03 - A*11 - A*30 - A*31 | | IEDB prediction tool |
| | 538-546 | CVNFNFNGL | | A*02 - A*68 | | IEDB prediction tool |
| | 583-592 | EILDITPCSF | | **A*26** | | 53 |
| **L4 pool (12 peptides)** | 588-597 | TPCSFGGVSV | | **B*07** | | 53 |
| | 598-608 | ITPGTNTSNQV | | A*68 - A*02 | | IEDB prediction tool |
| | 598-608 | ITPGTDTSNQV | | A*68 - A*02 | | IEDB prediction tool |
| | 660-670 | YECDIPIGAGI | | **B*40** | | 53 |
| | 733-741 | KTSVDCTMY | | A*30 - B*58 | | IEDB prediction tool |
| | 738-747 | CTMYICGDST | | A*68 | | IEDB prediction tool |
| | 747-756 | TECSNLLLQY | | **B*44** - **B*04** | * | 53 |
| | 756-765 | YGSFCTQLNR | | A*68 - A*11 - A*31 | | IEDB prediction tool |
| | 896-904 | IPFAMQMAY | | **B*35** - **B*51** - B*53 - A*30 | * | 53 |
| | 906-915 | FNGIGVTQNV | | A*68 | | IEDB prediction tool |
| | 908-917 | GIGVTQNVLY | | A*30 | | IEDB prediction tool |
| | 917-927 | YENQKLIANQF | | B*44 | | 53 |
| **L5 pool (12 peptides)** | 923-931 | IANQFNSAI | | **B*51** - B*35 - B*53 - B*58 - A*32 - A*68 | | 53 |
| | 958-966 | ALNTLVKQL | | **A*02** | | 30 |
| | 964-973 | KQLSSNFGAI | | A*02 - B*15 | | IEDB prediction tool |
| | 989-997 | AEVQIDRLI | | **B*44** - B*40 | * | 53 |
| | 996-1005 | LITGRLQSL | | **A*02** - B*08 | | 30 |
| | 1000-1008 | RLQSLQTYV | | **A*02** | | 30, 53 |
| | 1005-1014 | QTYVTQQLIR | | A*68 - A*11 - A*31 | | IEDB prediction tool |
| | 1016-1024 | AEIRASANL | | **B*44** - **B*40** | * | 53 |
| | 1029-1038 | MSECVLGQSK | | A*11 - A*68 | | IEDB prediction tool |
| | 1038-1047 | KRVDFCGKGY | | A*30 | | IEDB prediction tool |
| | 1048-1056 | HLMSFPQSA | | **A*02** | | 53 |
| | 1059-1067 | GVVFLHVTY | | **A*32** - B*15 - A*30 | | 53 |
| **L6 pool (10 peptides)** | 89-97 | GVYFASTEK | | **A*68** - **A*03** - **A*11** - A*30 | * | 31, 53 |
| | 95-104 | TEKSNIIRGW | | **B*44** | | 53 |
| | 162-170 | SANNCTFEY | | **B*35** - **A*29** - A*30 - A*01 - A*11 | * | 53 |
| | 162-170 | SANDCTFEY | | **B*35** - **A*29** | * | |
| | 169-177 | EYVSQPFLM | | **A*24** - A*23 | | 53 |
| | 202-210 | KIYSKHTPI | | **B*08** - A*32 - A*02 - A*30 | | 53 |
| | 224-231 | EPLVDLPI | | **B*51** - B*53 | | 53 |
| | 258-266 | WTAGAAAYY | | **B*35** - **A*26** - **A*29** | * | 53 |
| | 265-275 | YYVGYLQPRTF | | **A*24** - A*23 | | 53 |
| | 351-358 | YAWNRKRI | | **B*51** - B*08 - B*07 | | 53 |
| **L7 pool (10 peptides)** | 361-369 | CVADYSVLY | | **A*29** - **A*26** - B*35 - A*01 - A*11 - A*68 - A*30 | * | 53 |
| | 370-378 | NSASFSTFK | | **A*68** - A*11 - A*30 - A*03 | | 53 |
| | 378-386 | KCYGVSPTK | | **A*03** - A*30 | | 52, 53 |
| | 553-564 | TESNKKFLPFQQ | | **B*44** | | 53 |
| | 624-633 | IHADQLTPTW | | **B*53** - B*57 - B*58 | | 53 |
| | 628-636 | QLTPTWRVY | | **A*30** - B*15 | | 53 |
| | 634-643 | RVYSTGSNVF | | **B*15** - **A*24** - A*32 | * | 53 |
| | 637-646 | STGSNVFQTR | | **A*68** - **A*31** | * | 53 |
| | 725-733 | EILPVSMTK | | **A*03** - **A*68** - A*11 | * | 53 |
| | 779-789 | QEVFAQVKQIY | | **B*44** | * | 53 |
| **L8 pool (10 peptides)** | 821-829 | LLFNKVTLA | | **A*02** | | 30 |
| | 829-837 | ADAGFIKQY | | **B*44** | | 53 |
| | 857-865 | GLTVLPPLL | | **A*02** | | 53 |
| | 865-873 | LTDEMIAQY | | **A*01** - **B*35** - **A*29** | * | 31, 53, 56, 60 |
| | 869-877 | MIAQYTSAL | | **B*08** - A*02 - B*35 - A*68 - B*07 - B*15 | | 53 |
| | 1065-1073 | VTYVPAQEK | | **A*30** - **A*03** - A*11 - A*68 - A*31 | * | 53 |
| | 1093-1102 | GVFVSNGTHW | | **B*57** - B*58 | | 53 |
| | 1093-1102 | GVFVSDGTHW | | **B*57** | | |
| | 1099-1107 | GTHWFVTQR | | **A*11** - **A*31** - A*68 | | 31, 53 |
| | 1137-1146 | VYDPLQPELD | | **A*24** - A*23 | | 53 |
| **L9 pool (9 peptides)** | 1147-1155 | SFKEELDKY | | **A*29** | | 53 |
| | 1185-1193 | RLNEVAKNL | | **A*02** | | 30 |
| | 1192-1200 | NLNESLIDL | | **A*02** | | |
| | 1192-1200 | NLDESLIDL | | **A*02** | | |
| | 1201-1209 | QELGKYEQY | | **B*44** | * | 53 |
| | 1208-1216 | QYIKWPWYI | | **A*24** - **A*23** | * | 31, 52 |
| | 1220-1228 | FIAGLIAIV | | **A*02** - A*68 | | 30 |
| | 1256-1265 | FDEDDSEPVL | | B*40 | | 53 |
| | 1264-1272 | VLKGVKLHY | | **A*29** - B*15 - A*30 | | 53 |

**Hyper conserved SARS-Cov-2 spike sequences**

**Highly conserved SARS-Cov-2 spike sequences**

**Conserved SARS-Cov-2 spike sequences**

**C** **Conservation level**

**D** **Degenerated HLA recognition**

*In bold are the HLA restriction molecules identified experimentally; regular characters depict the HLA restriction alleles identified by prediction tools
